# Sphingolipid metabolism orchestrates the establishment of the adult hair follicle stem cell niche to control skin homeostasis

**DOI:** 10.1101/2024.01.09.574628

**Authors:** Franziska Peters, Susanne Brodesser, Kai Kruse, Hannes C.A. Drexler, Jiali Hu, Dominika Lukas, Esther von Stebut, Martin Krönke, Carien M. Niessen, Sara A. Wickström

## Abstract

Bioactive sphingolipids serve as an essential building block of membranes, forming a selective barrier that ensures subcellular compartmentalization and facilitates cell type-specific intercellular communication through regulation of the plasma membrane receptor repertoire. How cell type-specific lipid compositions are achieved and what is their functional significance in tissue morphogenesis and maintenance has remained unclear. Here, we identify a stem-cell specific role for ceramide synthase 4 (CerS4) in orchestrating fate decisions in skin epidermis. Deletion of CerS4 in the epidermis prevents the effective development of the adult hair follicle bulge stem cell (HFSCs) niche due to altered differentiation trajectories of HFSC precursors towards upper hair follicle and inner bulge fates. Mechanistically, HFSC differentiation defects arise from an imbalance of key ceramides and their derivate sphingolipids in HFSCs associated with hyperactivity of canonical Wnt signaling. Impaired HFSC niche establishment leads to disruption of hair follicle architecture and hair follicle barrier function, ultimately triggering a T helper cell 2 - dominated immune infiltration closely resembling human atopic dermatitis. This work uncovers a fundamental role for a cell state-specific sphingolipid profile in epidermal stem cell homeostasis and the role of an intact stem cell niche in maintaining an intact skin barrier.

## Introduction

The skin epidermis is a life-essential bi-directional barrier that protects animals from extrinsic insults including pathogens, toxins and irradiation, while preventing water loss from inside the body. Disruption of the barrier leads to a strong innate immune reaction and when unrepaired leads to chronic inflammatory skin diseases such as atopic dermatitis (AD) (Eyerich *et al*, 2015; Paternoster *et al*, 2015; Weidinger & Novak, 2016).

Given their function as a first line of defense, skin keratinocytes sustain constant damage, and in order to maintain tissue integrity these damaged cells are removed via terminal differentiation and subsequent tissue turnover rather than apoptosis (Kato *et al*, 2021). This homeostatic self-renewal process is driven by stem cells (Blanpain & Fuchs, 2009, 2014; Chacón-Martínez *et al*, 2018; Gonzales & Fuchs, 2017; Hsu *et al*, 2014; Schneider *et al*, 2009; Tumbar *et al*, 2004). The epidermis contains multiple stem cell populations that are required for compartment-specific tissue maintenance and repair. While the basal epidermal stem cells drive self-renewal of the interfollicular epidermis (IFE), the hair follicle and sebaceous glands are maintained by their own specific stem cells, the hair follicle stem cells (HFSCs), residing within the bulge stem cell niche (Blanpain & Fuchs, 2014; Chacon-Martinez *et al*, 2017; Hsu *et al*, 2011). While the key role of the bulge HFSCs in maintaining the renewal of the hair follicle and the hair shaft itself is well established, how this adult stem cell compartment becomes established during postnatal development is less clear.

The capacity of stem cells to self-renew or differentiate can be attributed to distinct metabolic states. Emerging evidence suggests that lipid metabolism plays a fundamental role in stem cell homeostasis (van Gastel *et al*, 2020) but the mechanisms are unclear. Ceramides, synthesized by ceramide synthases (CerS), are the essential building blocks of all cellular membranes and can also serve as signaling molecules. Ceramides have been implicated in regulating the balance between self-renewal and differentiation of neuronal stem cells, influencing signaling pathways that control cell fate decisions (Bieberich *et al*, 2003; He *et al*, 2014; Wang *et al*, 2008). CerS2-6 are expressed in mammalian skin (Levy & Futerman, 2010). CerS3 is essential for the formation of the functional epidermal barrier through the synthesis of extracellular μ-hydroxylated ultra-long chain ceramides, crucial components of the outer most layer of the skin, the stratum corneum (Eckl *et al*, 2013; Jennemann *et al*, 2012; Radner *et al*, 2013). CerS4 has been shown to regulate sebaceous gland homeostasis, hair follicle cycling, and homeostatic epidermal barrier function (Ebel *et al*, 2014; Peters *et al*, 2020; Peters *et al*, 2015). CerS4 has the highest substrate specificity towards long-chain ceramides, Cer18:0-22:0, key molecules in intracellular sphingolipid metabolism and essential building blocks of all cellular membranes (Tidhar *et al*, 2018). The mechanisms by which CerS4 regulates epidermal homeostasis and its role in stem cell regulation have remained unclear.

Here we show that CerS4 is crucial for sphingolipid metabolism required to direct stem cell lineage progression to establish the adult HFSC compartment. Deleting CerS4 in epidermal stem cells leads to defective establishment of the HFSC state, and these cells are inappropriately routed into upper hair follicle and inner bulge fates due to aberrant Wnt signaling. As a result, hair follicle architecture and barrier function are disrupted, leading ultimately to a T helper cell 2 (Th2) immune dominance with resemblance to AD. Collectively these data demonstrate a central role for lipid metabolism in tuning signaling required for stem cell fate regulation and the essential role of the HFSCs in maintaining the skin barrier.

## Results

### CerS4 is required for the establishment of the HFSC niche

Previous work had shown that constitutive deletion of CerS4 (CerS4^-/-^) in the entire animal and conditional deletion of CerS4 in the epidermis (from here on CerS4^epi-/-^) result in normal embryonic development of the epidermis but abnormal postnatal hair follicle cycling and progressive alopecia as well as impaired epidermal barrier function, acanthosis and hyperkeratosis (Ebel *et al*., 2014; Peters *et al*., 2020; Peters *et al*., 2015). To investigate the role of sphingolipid metabolism in controlling HFSC behavior, and to understand if HFSC intrinsic alterations in sphingolipid metabolism are associated with skin barrier disease, we carried out detailed analysis of the epidermis in mice where CerS4 was deleted in all epidermal stem cell compartments and their differentiated progeny using Keratin-14-Cre (Fig. 1A) (Hafner *et al*, 2004; Peters *et al*., 2015). Histological analyses in early postnatal mice revealed no obvious differences in the morphology of the interfollicular epidermis (IFE), but the pilosebaceous unit of these adolescent CerS4^epi-/-^ mice showed an altered structure characterized by an increased size of sebaceous glands already at postnatal day (P)19, indicating abnormalities in hair follicle morphology as an early event in the onset of skin pathology in these animals (Supplementary Fig. 1A).

**Figure 1.**
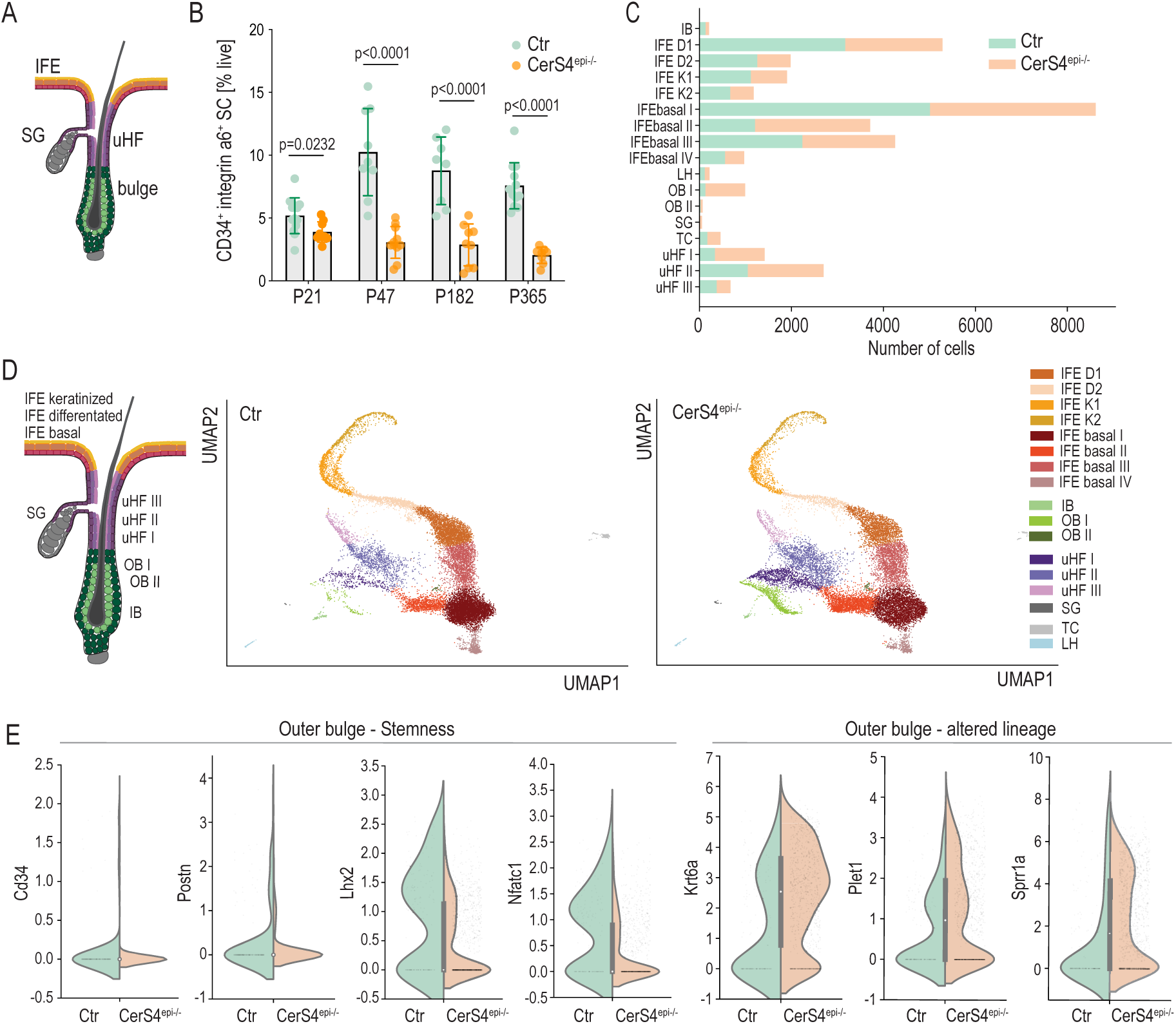
CerS4 is required for the establishment of the HFSC niche. **A.** Schematic illustration of the interfollicular epidermis (IFE) and the pilosebacous unit containing the sebaceous gland (SG) and the hair follicle (HF) and its bulge stem cell niche. Keratin-14-Cre is expressed in basal cells throughout the tissue, allowing Keratin-14-Cre-mediated deletion of CerS4 in all stem cell compartments. **B.** FACS-mediated quantification of CD34^+^ integrin α6^+^ HFSCs at P21 (n=13 Ctr, 9 CerS4^epi-/-^ mice), P47 (n=9 Ctr, 11 CerS4^epi-/-^ mice), P128 (n=8 Ctr, 9 CerS4^epi-/-^ mice), P365 (12 Ctr, 8 CerS4^epi-/-^ mice). Note progressive loss of stem cells in CerS4^epi-/-^ mice (mean± SD; unpaired t test). **C.** Cell counts of cell clusters determined by single cell RNA sequencing. **D.** Schematic of a resting phase hair follicle and UMAPs of cells identified using scRNAseq. Note comparable presence of all epidermal cell states but increased outer bulge population in CerS4^epi-/-^ mice (n=2 mice/genotype, pooled). **E.** Gene expression of indicated stem and progenitor cell marker genes in the outer bulge stem cell compartment in Ctr and CerS4^epi-/-^ skin.

To understand if the structural abnormalities of the pilosebaceous unit were associated with defects in the HFSC compartment in an epidermis-intrinsic manner, we quantified levels of CD34^+^ integrin-α6^+^ HFSCs from P21, P47, 6 mo and 1 y old control and CerS4^epi-/-^ mice. Interestingly CerS4^epi-/-^ mice showed reduced HFSC numbers already at P21 during postnatal hair morphogenesis (Fig. 1B), at the time when the stem cell niche is first established and when the CerS4^epi-/-^ mice still showed normal IFE but impaired hair follicle architecture.

To understand the role of CerS4 in the establishment of the HFSC compartment and in epidermal homeostasis more broadly, we isolated dorsal epidermis from P19 control and CerS4^epi-/-^ mice and generated single-cell transcriptome libraries (scRNAseq; BD Rhapsody). 2 mice/genotype were processed and sequenced. After individual quality control, all replicates were merged and analyzed together (17577 control cells / 17096 CerS4^epi-/-^ cells) to capture all cell populations. Cell types and states were identified based on cluster-specific gene expression and annotated based on previously published marker gene profiles (Joost *et al*, 2016). Our analyses revealed that the majority of the cells derived from the IFE. The IFE was divided in basal cells (IFE basal I-IV), differentiated (IFE D I-II) and keratinized (IFE K I-II) clusters. Furthermore, all major subsets of the hair follicle were detected and clustered. We detected two outer bulge stem cell compartments (OB I-II), the inner bulge (IB), and the upper hair follicle (uHF I-III) as well as sebocytes (SG). Further, Langerhans cells (LH) and T-cells (TC) formed the two non-epidermal clusters (Fig. 1C, D). Importantly, populations of the hair follicle showed alterations, characterized by a striking expansion of the outer bulge stem cell (OB I), the upper hair follicle (uHF I-II) and sebocyte (SG) populations in CerS4^epi-/-^ skin (Fig. 1C, D).

CerS4 transcripts were detected in the hair follicle and in sebaceous gland clusters, albeit with low levels as expected for an enzyme, as also validated by in situ RNA hybridization (Supplementary Fig. 1B, C). This hair follicle-specific expression pattern was confirmed by qRT-PCR analysis of FACS-purified (i) Sca1^-^ CD34^+^ integrin-a6^+^ HFSCs, (ii) Sca1^-^ CD34^-^ integrin-a6^+^ hair follicle progenitors and (iii) Sca1^+^ CD34^-^ integrin-a6^+^ IFE progenitors, where CerS4 mRNA expression was restricted to the HFSC compartment (Supplementary Fig. 1D).

As our data indicated high CerS4 mRNA expression in sebocytes, and a previous study had suggested that the hair follicle phenotype of CerS4-deficient mice results from altered sebum composition (Ebel *et al*., 2014), we asked if the observed hair follicle phenotypes were a result of altered sebaceous gland function in CerS4-deficient mice. To this end we deleted CerS4 specifically in the sebaceous glands using SCD3-Cre (Dahlhoff *et al*, 2016). Strikingly, mice lacking CerS4 in the sebaceous gland did not display any epidermal or hair follicle phenotypes (Supplementary Fig. 1E). There were no macroscopic signs of alopecia, and careful histological analysis of skin sections showed no alterations in hair follicle morphology nor structural abnormalities of the IFE (Supplementary Fig. 1F, G). Finally, flow cytometry analyses of freshly isolated epidermal cells from P47, P58 and 6 mo old control and CerS4^SCD3-/+^ mice showed comparable levels of CD34^+^ integrin-α6^+^ HFSCs (Supplementary Fig. 1I). The data demonstrate that CerS4 function in sebocytes is not required to establish the HFSC niche or to control structural integrity of the hair follicle, the sebaceous gland, or IFE.

Having excluded a causative role of CerS4 in the sebaceous gland in the development of the hair follicle phenotype, we focused our attention to the HFSC compartment, in particular on the observed expansion of the outer bulge stem cell population (OB I) in the CerS4^epi-/-^ mice. Gene expression profiling of key hair follicle outer bulge stem cell markers and transcriptional master regulators such as *Cd34* and *Postn* revealed that their expression was slightly reduced in CerS4-deficient outer bulge cells. In particular, levels of *Lhx2,* essential for maintaining HFSC quiescent and ensuring linage stability (Folgueras *et al*, 2013) and of the master regulator of quiescent HFSCs, *Nfatc1* (Horsley *et al*, 2008), were substantially reduced (Fig. 1E). Strikingly, markers of differentiation into inner bulge-like and upper hair follicle states, such as *Krt6a, Plet1* and *Sprr1* were increased in this compartment (Fig. 1E), indicative of altered lineage identity of the outer bulge cells.

Collective, these data show that CerS4 in HFSCs is essential to establish the adult stem cell compartment and to assure lineage fidelity. This altered lineage identity likely explains the shifts in population ratios and drives progressive loss of the HFSC compartment, resulting in alopecia.

### CerS4 controls hair follicle stem cell lineage specification

As our data pointed to altered lineage identity and population proportions of the early HFSCs, we sought to predict lineage trajectories of these cells from the scRNAseq data using Slingshot (Street *et al*, 2018). Ordering cells isolated from control skin along a path starting from the outer bulge stem cells population revealed a trajectory from outer bulge stem cells to inner bulge stem cells, and a path from outer bulge stem cells to the upper hair follicle clusters. Cell lineage inference of CerS4^epi-/-^ skin indicated an additional, abnormal trajectory from the outer bulge hair follicle stem cells to the sebaceous cluster (Fig. 2A). Pseudotime analyses further confirmed altered linage progression, when focusing on a trajectory along upper hair follicle cells, outer bulge and inner bulge stem cells in CerS4^epi-/-^ mice. Here, the upper hair follicle marker Plet1 remained high in the outer bulge stem cells, whereas expression of the outer bulge stem cell factor Lhx2 was not turned on efficiently. In contrast, the outer bulge cells prematurely upregulated the inner bulge marker Krt6 (Fig. 2B).

**Figure 2.**
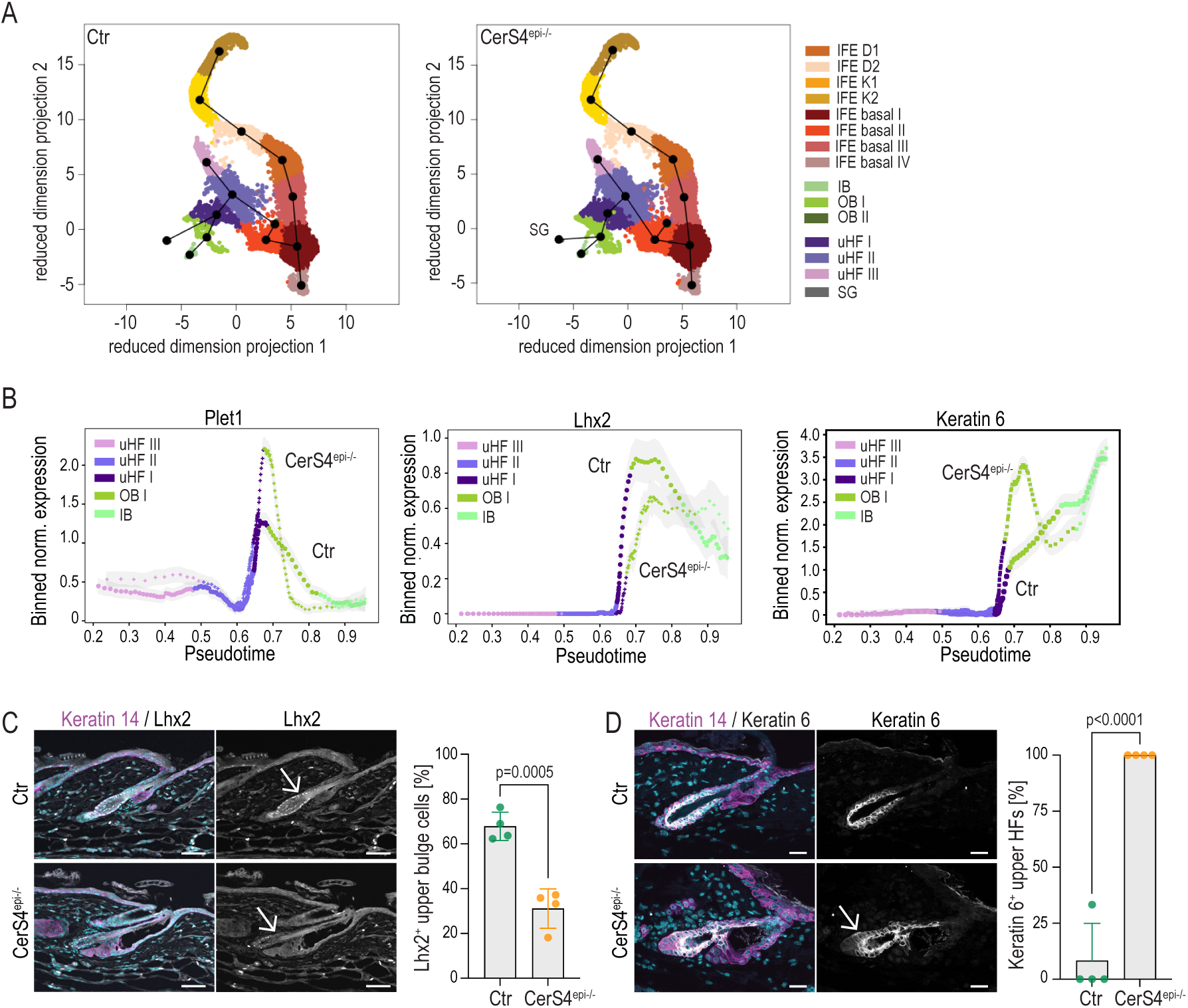
CerS4 controls hair follicle stem cell lineage specification. **A.** Linage trajectories as determined by SlingShot of control (Ctr) and CerS4^epi-/-^ cells from scRNAseq data (n=2 mice/genotype, pooled). Note abnormal trajectory from outer bulge cells into sebaceous gland. **B.** Pseudotime analyses of Ctr and CerS4^epi-/-^ cells from scRNAseq data (n=2 mice/genotype, pooled) confirming altered linage progression with expression of Plet1 remaining high, Lhx2 low and Krt6 high in CerS4^epi-/-^ outer bulge stem cells. **C.** Representative images and quantification of P21 back skin sections stained for Lhx2 (grey) and Keratin-14 (magenta) show reduced expression of Lhx2 in bulge stem cells (Scale bars 50µm; n=4 mice/genotype; mean± SD; unpaired t test). **D.** Representative images and quantification of P21 control and CerS4^epi-/-^ back skin sections stained for Keratin-6 (grey), Keratin-14 (magenta) show ectopic Keratin-6 expression in the outer bulge and upper hair follicle of CerS4^epi-/-^ mice. (Scale bars 25µm; n=4 mice/genotype; mean± SD; unpaired t test).

As the data so far indicated defective commitment of outer bulge cells that instead displayed features of inner bulge-like and upper hair follicle lineages, we sought to further investigate where this altered differentiation occurs. Immunofluorescence analyses of back skin sections from P21 mice showed decreased *Lhx2* expression in the CerS4^epi-/-^ outer bulge stem cell region of CerS4^epi-/-^mice, most prominently visible in the upper part of the bulge (Fig. 2C). Lhx2 loss has been reported to lead to a failure to maintain HFSC quiescence and a progressive transformation of the niche into a sebaceous gland (Folgueras *et al*., 2013).

Interestingly, a reduction in the pearl string-like cell composition in the upper bulge and an increased cell size resembling a sebocyte-like cell shape was observed in CerS4^epi-/-^ back skin (Fig. 2C, D). Furthermore, we observed expansion of the inner bulge identity marker Krt6 protein expression into outer bulge stem cells and along the infundibulum in CerS4^epi-/-^ hair follicles, whereas in control mice Krt6 was restricted to the inner bulge (Fig. 2C).

Collectively, these data indicate that CerS4-deficiency triggers an abnormal lineage trajectory of outer bulge stem cells into upper and inner hair follicle as well as sebaceous gland-like fates, in providing a likely mechanism for the inefficient establishment and further gradual depletion of the quiescent HFSC compartment.

### CerS4 regulates HFSC differentiation in a stem cell autonomous manner

To gain insight into the cellular and molecular mechanisms by which CerS4 cell-autonomously regulates HFSC lineage trajectories, we utilized an *ex vivo* organoid culture system that promotes the growth of CD34^+^ integrin α6^+^ and CD34^-^ integrin-α6^+^ cells, which based on extensive transcriptome and marker expression analyses represent a mixture of HFSCs, hair follicle outer root sheath (ORS) cells and inner bulge cells (collectively termed non-HFSCs). Sebocytes, cells from keratinized layers of the epidermis, or cells of non-keratinocyte lineages are not detected in these organoids (Chacon-Martinez *et al*., 2017; Kim *et al*, 2020). We generated organoids from freshly isolated P19 control and CerS4^epi-/-^ epidermis and confirmed that CerS4 expression was maintained in both HFSC (CD34^+^ integrin a6^+^) and non-HFSC (CD34^-^ integrin a6^+^) populations in control organoids (Fig. 3A, B). Strikingly, CerS4^epi-/-^ organoids showed altered morphology characterized by smaller size and loss of cohesion of peripheral cells from the organoid clusters (Fig. 3C), as well as strongly attenuated growth and impaired long-term maintenance within a 14d culture period (Supplementary Fig. 2A). CerS4^epi-/-^ organoids showed reduced HFSC numbers (Fig. 3D).

**Figure 3.**
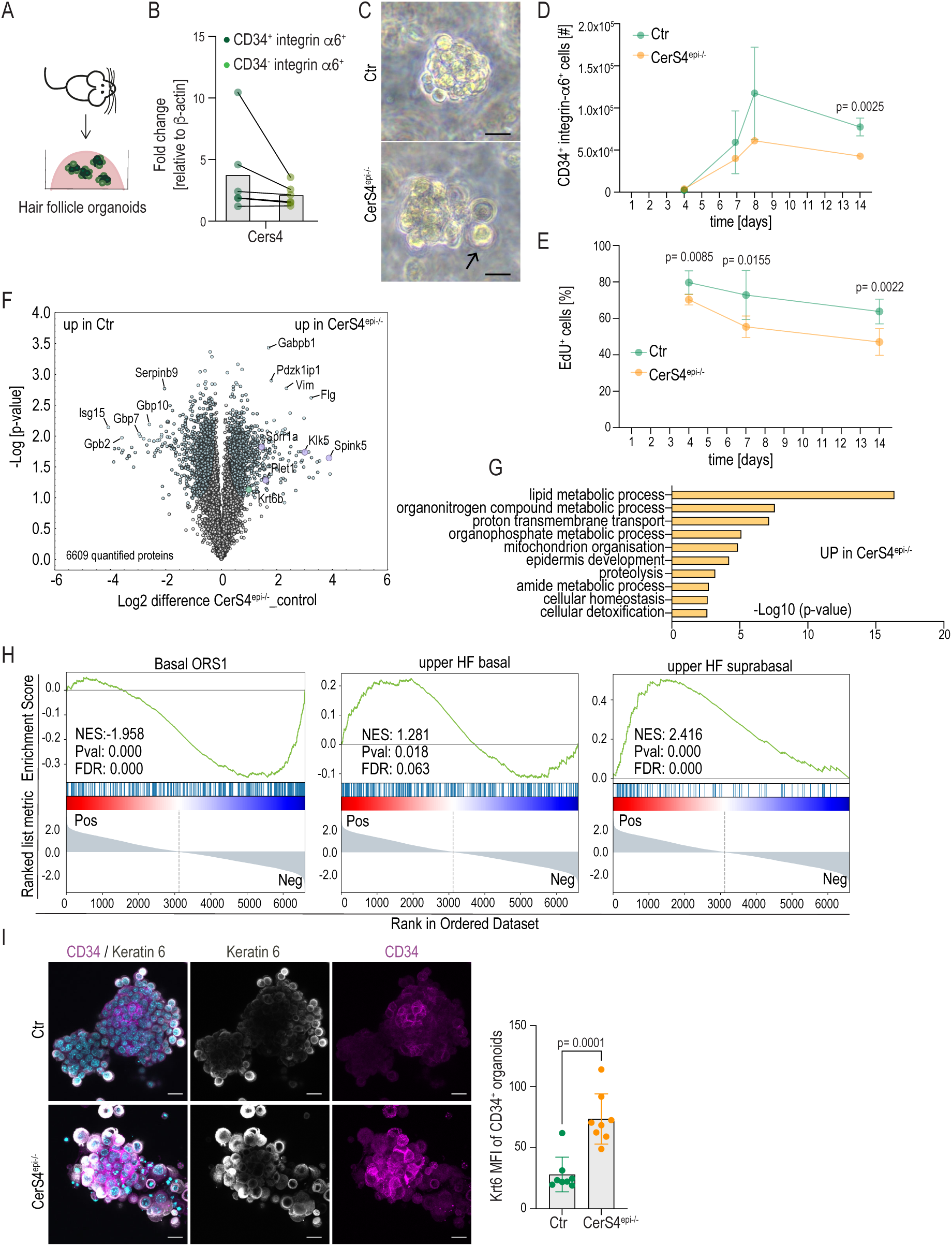
CerS4 regulates HFSC differentiation in a stem cell autonomous manner. **A.** Schematics of HFCS organoid generation. **B.** qRT-PCR analyses of Cers4 expression of FACS purified CD34^+^ α6^+^ integrin HFSCs and CD34^-^ integrin α6^+^ non-HFSCs from organoids (n=5 independent experiments). **C.** Representative brightfield images of control (Ctr) and CerS4^epi-/-^ organoids cultured for 4 days. Arrows mark non-cohesive peripheral cells in CerS4^epi-/-^ organoids. Scale bar 10µm. **D**. FACS-based quantification CD34^+^ integrin α6^+^ HFSCs from organoids cultured for time points indicated show reduced HFSCs in CerS4^epi-/-^organoids (n=4 mice/genotype for 14d; 6 mice/genotype all other time points; mean± SD; unpaired t-test). **E.** Quantification of EdU-positive cells after a 24h chase of control and CerS4^epi-/-^ organoids cultured for indicated time points (n=6 mice/genotype; mean± SD; unpaired t-test). **F.** Volcano plot of differentially expressed proteins in control and CerS4^epi-/-^ organoids (n=4 mice/genotype). **G.** GO-term enrichment analyses of differentially expressed proteins in CerS4^epi-/-^ organoids. **H.** GSEA of the differential protein expression of control and CerS4^epi-/-^ organoids indicate under-representation of proteins from gene expression signatures from outer bulge and over-representation of proteins from upper hair follicle signatures (from Joost et al., 2016) in CerS4^epi-/-^ organoids. **I**. Representative images and quantification of control and CerS4^epi-/-^ organoids stained for Keratin-6 (grey) and Keratin-14 (magenta) show increased Keratin-6 and abundance of loosely attached, differentiating cells in CerS4^epi-/-^ (Scale bar 20µm; n=8 organoids/genotype; mean± SD; unpaired t-test).

The non-HFSC population was also slightly but not statistically significantly reduced (Supplementary Fig. 2B). No increase in apoptosis at d4 or d7 and only a minor increase at d14 of culture was detected in CerS4^epi-/-^ organoids (Supplementary Fig. 2C). In contrast, EdU incorporation experiments showed reduced proliferation of CerS4^epi-/-^ organoids already at d4 (Fig. 3E, Supplementary Fig. 2D, E). Overall, in both control and CerS4^epi-/-^ organoids the CD34^+^ integrin α6^+^ HFSCs showed higher rates of proliferation than the CD34^-^ integrin a6^+^ progenitors indicating that HFSCs are the main drivers of organoid growth.

To understand the mechanisms of reduced HFSC numbers and to mechanistically link the organoid phenotypes to the *in vivo* observations of altered differentiation trajectories, we performed quantitative proteomics on d8 control and CerS4^epi-/-^ organoids. A total of 6609 proteins could be quantified across all replicates, of which 399 (FDR q<0.05, s0=0.01, fold change> 2) were differentially expressed (Fig. 3F; Supplementary Table 1). GO Biological Processes (GOBP) term analysis implicated metabolic processes, epidermal development and inflammation as most strongly altered, with no major differences in proteins associated with cell death and growth (Fig. 3G, Supplementary Fig. 2F). Importantly, proteins with increased expression in CerS4^epi-/-^ organoids included Krt6 and Plet1, key markers of the differentiated inner bulge/companion layer and the upper hair follicle, respectively, that were also increased in the outer bulge population *in vivo*. Other companion layer/upper hair follicle marker proteins were also upregulated, including Klk5, Spink5, Sprr5 and Filaggrin (Furio *et al*, 2015; Yang *et al*, 2004; Ziegler *et al*, 2019), pointing to a cell-autonomous nature of the HFSCs differentiation defect in CerS4^epi-/-^ HFSCs. The most downregulated proteins included Serpinb9, Isg15, Gbp2, Gbp7 and Gbp10, known to be related to immune responses (Bird *et al*, 2014; Perng & Lenschow, 2018; Tretina *et al*, 2019), indicating that loss of stem cells was associated with attenuation of keratinocyte intrinsic immune regulation.

To investigate if these differential protein expression patterns would indeed reflect differences in HFSC differentiation trajectories in CerS4^epi-/-^ HFSCs, we systematically compared the differences in protein expression to known markers of distinct progenitor cell linages. Indeed, Gene Set Enrichment Analysis (GSEA; (Subramanian *et al*, 2005) revealed enrichment of upper hair follicle markers as well as the sebaceous gland in the upregulated proteome of CerS4^epi-/-^ organoids along with negative enrichment of basal outer root sheath/stem cell compartment markers (Fig. 3H; Supplementary Fig. 2G).

Consistent with the data obtained from proteomics, immunofluorescence analyses of control and CerS4^epi-/-^ organoids revealed increased levels of Krt6^+^cells in CerS4^epi-/-^ organoids (Fig. 3I). An increase in Krt6^+^ cells was also found in the periphery of the organoid clusters surrounding the CD34^+^ HFSCs, explaining the differences seen in organoid morphology with brightfield imaging (Fig. 3C). These loosely attached differentiating cells do not contribute to proliferation in the organoids, explaining the difference in cell numbers (Supplementary Fig. 2A).

Collectively these data indicated reduced HFSC maintenance and increased differentiation of cells into an upper hair follicle and inner bulge/companion layer states in the organoids, confirming that CerS4 activity in HFSCs is required for maintenance of a stable, undifferentiated HFSC population.

### CerS4 activity is required to maintain membrane lipid homeostasis

The mechanisms by which ceramide synthases regulate lipid metabolism are highly complex due to an intricate interregulation of sphingolipid metabolism through these enzymes (Mullen *et al*, 2011) (Fig. 4A). Given the critical roles of sphingolipids in membrane composition in intracellular compartments and the plasma membrane, we hypothesized that CerS4 is required to establish a specific membrane topology to orchestrate signaling in HFSCs. To test this, we performed quantitative lipidomics analyses of control and CerS4^epi-/-^ organoids to quantify sphingolipids.

**Figure 4.**
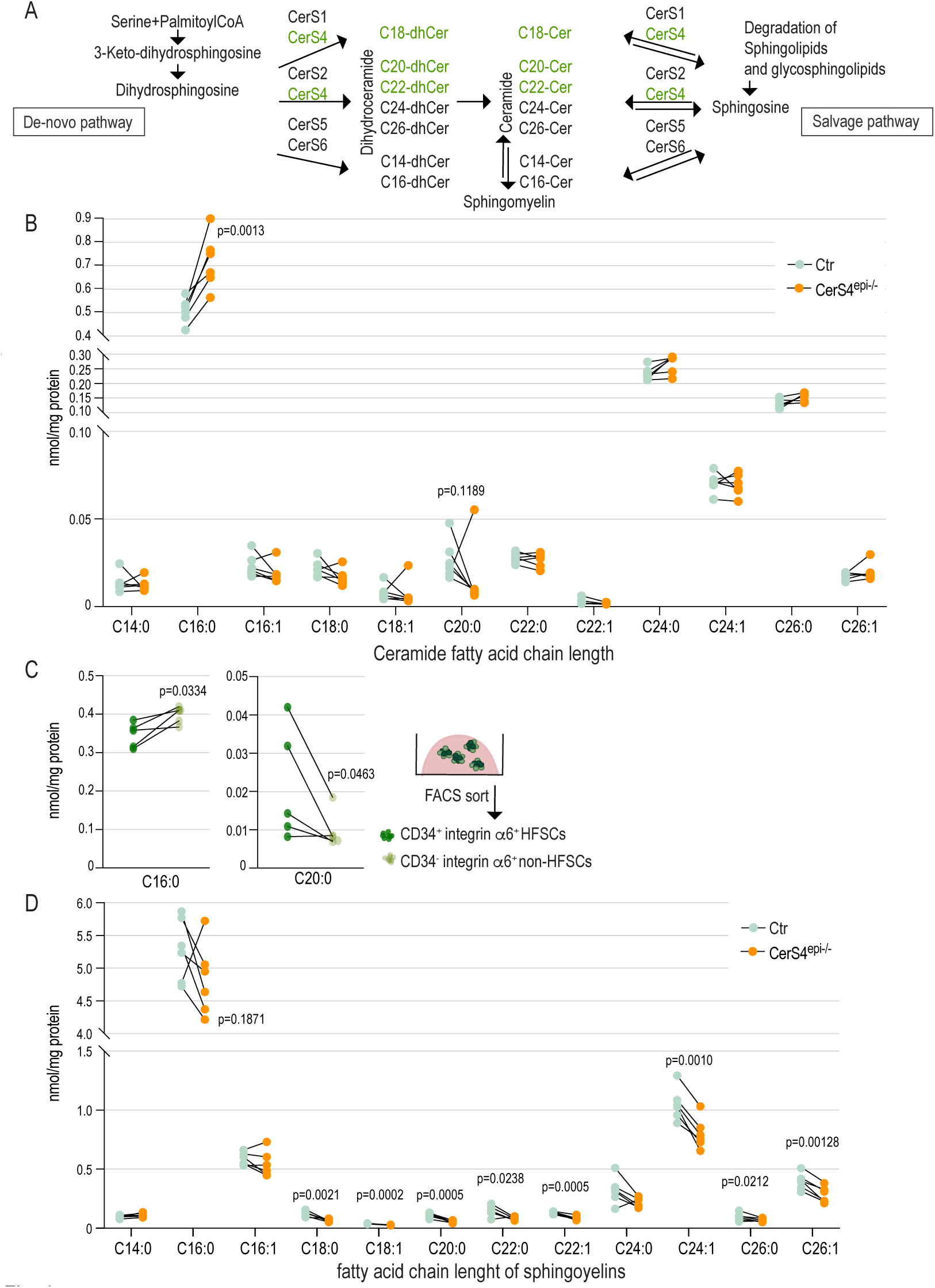
CerS4 activity is required to maintain membrane lipid homeostasis. **A.** Schematic of the substrate specificity of ceramide synthases (CerS) **B.** Ceramide levels in control and CerS4^epi-/-^ organoids determined by quantitative LC-ESI-MS/MS analysis. Note increased C16:0 and slightly decreased C18:0 and C20:0 ceramides (n=6 mice/genotype; ratio-paired t-test). **C.** Levels of C16:0 and C20:0 ceramides in FACS purified CD34^+^ integrin α6^+^ HFSCs and CD34^-^ integrin α6^+^ non-HFSCs determined by quantitative LC-ESI-MS/MS analysis (n=5 mice/genotype; ratio-paired t-test). **D.** Sphingomyelin levels in control and CerS4^epi-/-^ organoids determined by quantitative LC-ESI-MS/MS analysis. Note decreased C18:0, C18:1, C20:0, C22:0, C22:1, C24:1, C26:0 and C26:1 sphingomyelins (n=6 mice/genotype; ratio-paired t-test).

Analysis of sphingolipids showed an increase in ceramide C16:0 in CerS4^epi-/-^ organoids (Fig. 4B). A similar increase was observed in FACS purified wild-type non-HFSC cells compared to wild-type HFSCs (Fig. 4C), linking ceramide C16:0 to stem cell differentiation. Further, a moderate albeit statistically not significant decrease in ceramides with a fatty acid chain length of C18-C22 reflecting CerS4’ substrate preferences, were detected in CerS4^epi-/-^ organoids (Fig. 4B). A decrease in ceramide C20:0 was also seen in purified wild type non-HFCSs compared to HFSCs (Fig. 4C), linking this fatty acid to the stem cell state. Ceramides with a fatty acid chain length of C14:0, C24:0, C24:1, C26:0 and C26:1 were largely unaltered (Fig. 4B, Supplementary Fig. 3).

Ceramides serve as the starting point for the synthesis of sphingomyelin that plays a crucial role in maintaining the structural integrity and fluidity of cell membranes for proper signaling. Importantly, analysis of sphingomyelin levels revealed a decrease in sphingomyelins containing C18:0, C18:1, C20:0, C22:0, C22:1, C24:1, C26:0, C26:1 fatty acids, indicative of altered membrane composition in CerS4^epi-/-^ organoids (Fig. 4D). Thus, loss of CerS4 results in a decrease in specific sphingomyelin levels likely due to the reduction of its direct substrates, the C18:0, C18:1, C20:0, C22:0, C22:1 ceramides and compensatory dysregulation in other species. As ceramide and sphingomyelin cycling are essential for proper cell signaling the imbalance in ceramide and sphingomyelin levels likely impacts on transmembrane signaling.

### Altered response to Wnt signaling drives abnormal stem cell states

To investigate potential transmembrane signaling defects by which CerS4-dependent lipid homeostasis could control HFSC maintenance and differentiation, we returned to the *in vivo* scRNAseq data and focused on cell-cell communication that is known to be critical for determining HFSC state and behavior (Hsu *et al*., 2011). Using receptor-ligand interaction analyses (CellPhoneDB; (Efremova *et al*, 2020), we analyzed cellular communication within the outer bulge cells themselves as well as between the outer bulge and inner bulge. The most prominently implicated changes in CerS4^epi-/-^ mice were an increase in canonical and non-canonical Wnt as well as in BMP signaling (Fig. 5A; Supplementary Fig.4A). This was intriguing as Wnt and BMP are key signaling pathways controlling HFSC state establishment and maintenance (Daszczuk *et al*, 2020).

**Figure 5.**
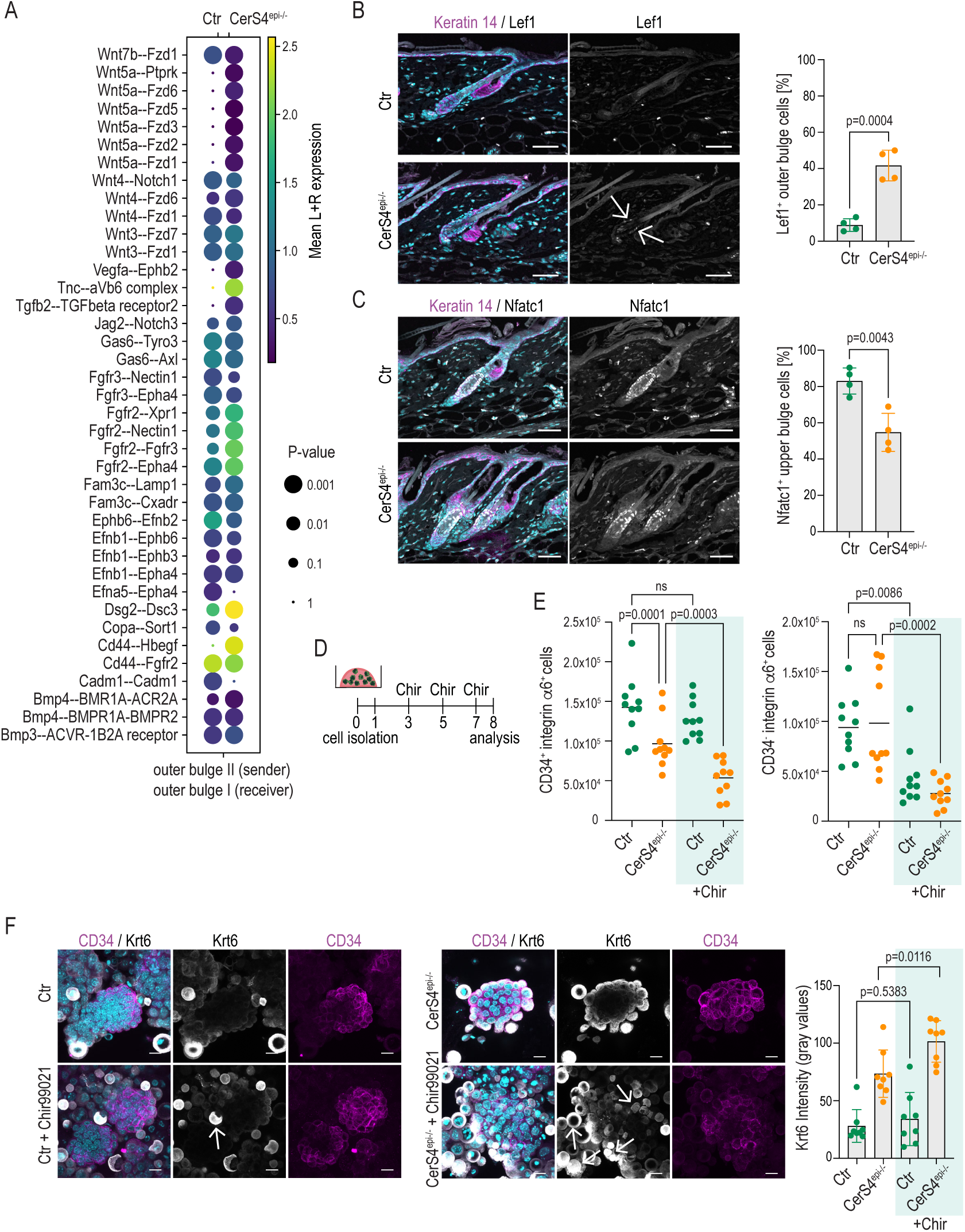
Altered response to Wnt signaling drives abnormal stem cell states. **A.** Receptor-ligand interaction analyses from scRNAseq data of control (Ctr) and CerS4^epi-/-^ skin. Communication within outer bulge cells is illustrated. Note altered Wnt signaling in CerS4^epi-/-^ outer bulge. **B**. Representative images and quantification of P21 Ctr and CerS4^epi-/-^ back skin sections stained for Lef1 (grey) and Keratin 14 (magenta). Note increased Lef1 expression in CerS4^epi-/-^ outer bulge (Scale bar 50µm; n=4; mean± SD; unpaired t-test). **C.** Representative images and quantification of P21 back skin sections stained for Nfatc1 (grey) and Keratin-14 (magenta). Note decreased levels of Nfatc1 in CerS4^epi-/-^ upper bulge (Scale bar 50µm; n=4; mean± SD; unpaired t-test). **D.** Experimental outline for Wnt-activation using Chir99021. **E.** FACS- based quantification of HFSCs and non-HFSCs from Ctr and CerS4^epi-/-^ organoids treated with Chir99021. Note decrease in HFSCs in Chir99021-treated CerS4^epi-/-^ organoids but not in control organoids (n=10 mice/genotype, mean± SD; unpaired t-test). **F** Representative images and quantification of Ctr and CerS4^epi-/-^ organoids treated with Chir99021 and stained for Keratin-6 (grey) and Keratin-14 (magenta). Note increased Keratin-6 expression and loss of cell cohesion in CerS4^epi-/-^ organoids (Scale bar 20µm; n=8 organoids/genotype; mean± SD; unpaired t-test).

Wnt signaling is particularly critical for the establishment of the adult HFSC population, in which Wnt activity is required to initially establish the hair follicle lineage but then its activity needs to attenuate to establish HFSCs and prevent their differentiation (Xu *et al*, 2015). We thus asked if the CerS4-deficient HFSCs were unable to enter this required “Wnt-low” state upon initial specification of the hair follicle lineage. Indeed, scRNAseq data showed increased expression of the Wnt target genes Lef1 and Tcf4 in CerS4^epi-/-^ outer bulge cells (Supplementary Fig. 4B). Immunofluorescence analysis of back skin confirmed an increase in Lef1 protein levels in CerS4^epi-/-^ outer bulge stem cells *in vivo* (Fig. 5B). In addition, *Nfatc1* protein expression was decreased in the upper bulge of CerS4^epi-/-^ hair follicles (Fig. 5C). BMP signaling activates *Nfatc1* expression in the bulge to maintains bulge stem cells in a quiescent state (Horsley *et al*., 2008), further indicating increased activation of CerS4^epi-/-^ HFSC. This data suggested that CerS4-deficient outer bulge cells are unable to enter a quiescent, Wnt-low state.

To directly test Wnt responsiveness of CerS4-deficient cells, we increased Wnt activity in HFSC organoids by treating them with Chir99021, a small molecule inhibitor of the Wnt inhibitor glycogen synthase kinase 3 (GSK-3) (Fig. 5D). Flow cytometry-mediated quantification of HFSC (CD34^+^/integrin a6^+^) and progenitor (CD34^-^ /integrin a6^+^) numbers in control organoids revealed no change in HFSCs but a clear decrease in progenitor cell numbers triggered by high Wnt (Fig. 5E). These data indicate that these control progenitors but not HFSCs respond to increased Wnt by increased differentiation and associated cell cycle exit. Strikingly, the number of CerS4-deficient stem cells was strongly reduced upon activation of Wnt, while the progenitor response was similar to controls (Fig. 5E), showing that CerS4-deficient HFSCs are sensitive to high Wnt conditions, suggesting that CerS4- deficient HFSC are in a more “primed” state towards differentiation.

Immunofluorescence analyses revealed an Wnt-induced increase in Sox9 expression in CD34hi control HFSCs, as expected (Supplementary Fig. 4C). Consistent with their higher sensitivity to Wnt, CerS4-deficient HFSCs responded to the increased Wnt with a stronger increase in Sox9 expression (Supplementary Fig. 4C). Further, immunofluorescence analysis of control organoids subjected to Wnt activation showed an increase in the differentiated Krt6^+^ cells loosely associated with the periphery of the CD34^+^ HFSC cluster, whereas the CD34^+^ HFSC core remained compact (Fig. 5F). In striking contrast, CerS4^epi-/-^ organoids showed a strong decrease in compactness and increased Krt6 expression throughout the organoid (Fig. 5F), explaining the differences seen in HFSC (CD34^+^/integrin a6^+^) and progenitor (CD34^-^ /integrin a6^+^) numbers in flow cytometry-mediated quantification (Fig. 5E). Collectively this data shows that CerS4^epi-/-^ HFSCs are sensitized to Wnt signals, leading to loss of this population, thus explaining the observed inefficient establishment and gradual depletion of bulge HFSCs *ex vivo* and in CerS4-deficient mice.

### Deletion of Cers4 leads to inflammatory alterations resembling human atopic dermatitis

CerS4-deletion in mice showed macroscopic and histological characteristics of an AD-like eczema (Fig.1B,C; (Peters *et al*., 2020)). We next tested whether this phenotype may be secondary to CerS4 deficiency related impaired HFSC dynamics. AD is associated with immune dysregulation, particularly a Th2 dominance (Weidinger *et al*, 2018). Innate lymphoid cells type 2 (ILC2) represent a subset of innate lymphoid cells (Neill *et al*, 2010) that can promote allergic inflammation at multiple barrier organs (Vivier *et al*, 2018). ILC2 can be activated by signals from the epidermis to promote the differentiation of type 2 helper T-cell (Th2) cells, eventually amplifying Th2 responses (Salimi *et al*, 2013). Aberrant HFSC niche establishment impairs epidermal tissue architecture, which likely impacts on ILC2 immune niche establishment. Hence, we investigated immune cell populations in CerS4^epi-/-^ skin at the time point when hair follicle abnormalities were present but IFE architecture were only beginning to emerge.

Immunofluorescence analyses of back skin sections from P19, P21 and P58 mice showed no difference in CD45^+^ leucocyte numbers and localization when comparing the CerS4^epi-/-^ to control skin, indicating no difference in overall immune cell numbers in CerS4^epi-/-^ skin (Supplementary Fig. 5A). Consistently, FACS analyses revealed no differences in CD45^+^ leucocytes in CerS4^epi-/-^ mice at P44. To be able to determine immune cell subsets based on their surface protein expression., the B- and T cell lineage was stained using antibodies against CD3^+^, CD19^+^, TCRβ^+^, TCRγο^+^, CD5^+^, FCeR1^+^ and gated as linage positive or negative. Analysis of immune cell subsets showed a slight increase in Th2 cells (viable CD45^+^ cells, linage^+^, CD90.2^+^, CD4^+^) and an increase in activated ILC2 (viable CD45^+^ cells, lineage^-^, CD90.2^+^, CD25^+^) in CerS4^epi-/-^ skin (Fig. 6A). No alterations in non-activated GATA3^+^ ILC2 (viable CD45^+^ cells, lineage^-^, CD90.2^+^, Gata3^+^), ILC1 (viable CD45^+^ cells, linage^-^, Nkp46^+^, CD11b^-^, Rorγt^-^, Eomes^-^) and natural killer (NK) cells (viable CD45^+^ cells, linage^-^, Nkp46^+^, CD11b^-^, Rorγt^-^, Eomes^+^) were observed. The data indicate a shift towards a Th2 dominance in CerS4^epi-/-^ mice, associated with activated CD25^+^ ILC2.

**Figure 6.**
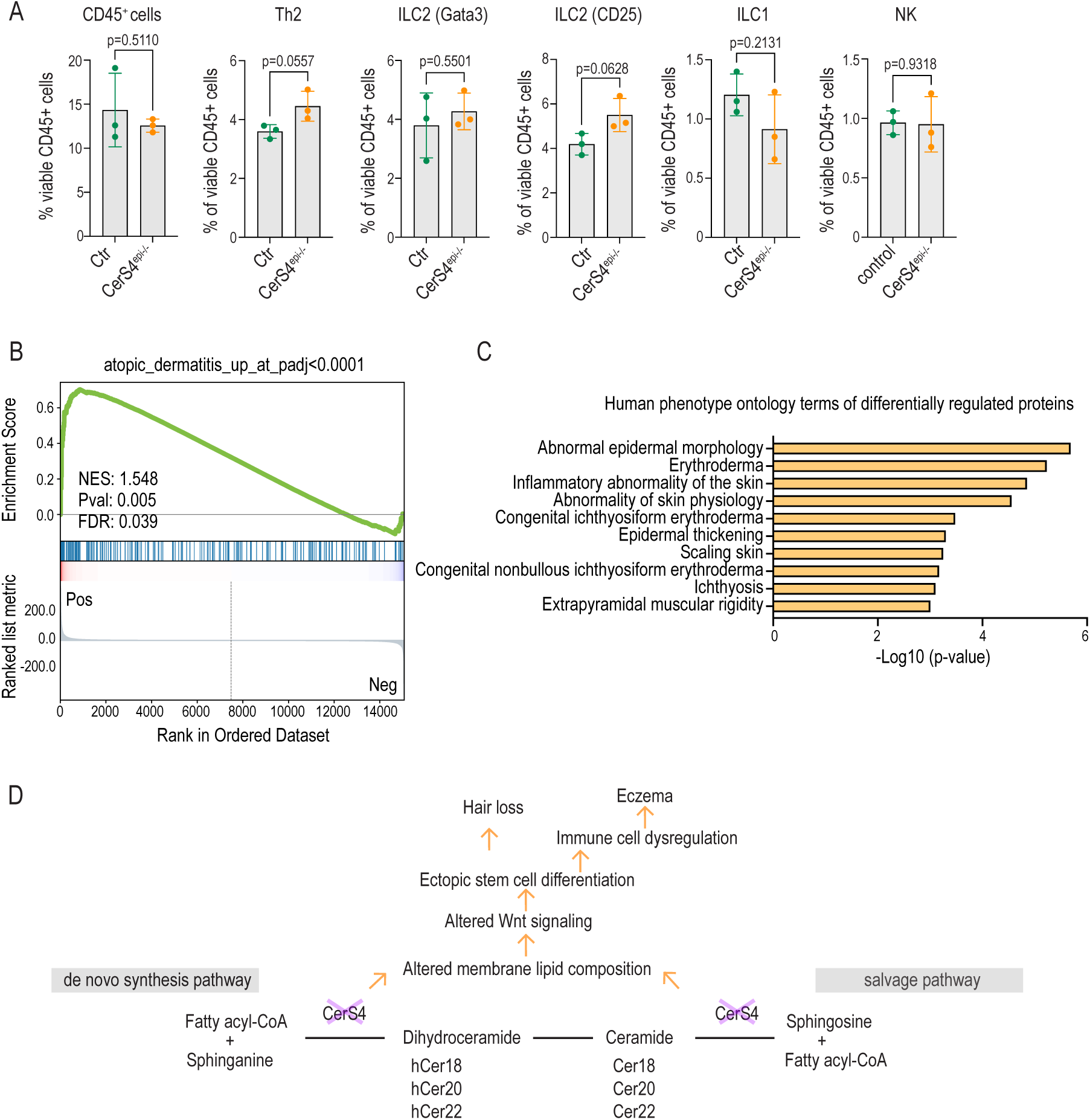
Deletion of Cers4 leads to inflammatory alterations resembling human atopic dermatitis. **A.** FACS analysis of CD45, Th2, ILC2(Gata3), ILC2(CD25), ILC1 and NK cells in control and CerS4^epi-/-^ ears (n=3 mice/genotype, mean± SD; unpaired t-test). **B.** GSEA of transcriptional profiles from patients diagnosed with atopic dermatitis (Federico *et al*., 2020) in differentially expressed genes in control and CerS4^epi-/-^ skin. **C.** Annotation enrichment analysis of in human phenotype ontology terms in differentially expressed proteins in control and CerS4^epi-/-^ organoids. **D.** Model of how CerS4-deletion in HFSCs leads to hair loss and eczema.

This AD-like inflammatory profile prompted us to investigate if CerS4-deletion would trigger gene expression alterations that share features found in human AD. To this end we compared data from CerS4^epi-/-^ skin with ortholog transcriptional profiles of 20 patients diagnosed with AD (Federico *et al*, 2020). GSEA of the health versus involved human skin compared to CerS4^epi-/-^ skin revealed an overlapping transcriptional profile of involved skin of patients with AD and CerS4^epi-/-^ skin (Fig. 6C). The data indicates that CerS4^epi-/-^ mice *in vivo* mimic transcriptional characteristics of human AD. Finally, we asked if these changes are related to altered HFSC function. Mining the Human Phenotype Ontology database with the differential protein expression dataset from control and CerS4^epi-/-^ organoids revealed erythroderma, epidermal acanthosis, abnormal epidermal morphology and inflammatory abnormality of the skin as the most prominent hits (Fig. 6B, Supplementary Table 2), where erythroderma describes exacerbation of an underlying skin disease, such as AD. Collectively this data indicate that cell-autonomous defects in HFSC homeostasis induced by loss of CerS4 trigger a transcriptional signature, a HFSC-intrinsic protein expression profile and a Th2 dominated immune phenotype, that shares features with human AD.

## Discussion

Emerging evidence suggests that lipid metabolism plays a fundamental role in stem cell homeostasis (van Gastel *et al*., 2020) but the mechanisms remain unclear. Our study shows that a specific ceramide profile is required for the establishment of an adult stem cell niche, the bulge HFSCs. Deletion of the ceramide synthase, CerS4 that is specifically expressed by the hair follicle cells led to misregulation of cell fate trajectories, where cells were routed towards upper hair follicle and inner bulge identities instead of HFSCs. Mechanistically, the hair follicle cells show altered ceramide and sphingomyelin profiles, associated with aberrant Wnt signaling. The inability to establish the HFSC niche leads to progressive alopecia and strikingly, a transcriptional signature, a keratinocyte intrinsic protein expression profile and a Th2 dominated immune phenotype that shares features of human AD.

The finding that CerS4 expression was restricted to the hair follicle where it regulated a specific cell fate transition of HFSCs, indicates that stem cells require establishment and dynamic maintenance of a specific CerS4-dependent lipid profile to facilitate cell type specific signaling. Our *in vivo* and organoid data point to a cell autonomous defect in Wnt signaling as the mechanism for altered stem cell differentiation. Wnt signaling is essential for establishing the hair follicle compartment and for proper regulation of HFSCs. The canonical Wnt pathway involves the stabilization and nuclear translocation of β-catenin. In the nucleus, β-catenin associates with transcription factors of the T-cell factor/lymphoid enhancer factor (TCF/LEF) family, leading to the transcriptional activation of i.e. c-Myc, Cyclin D1 or Axin2. Wnt ligands are secreted proteins that bind to cell surface receptors, initiating intracellular signaling cascades. Receptors involved in Wnt signaling include Frizzled receptors and low-density lipoprotein receptor-related proteins (LRP). The proper regulation of Wnt signaling is crucial for maintaining the balance between stem cell proliferation and differentiation. Wnt signaling is implicated in determining the fate of epidermal stem cells, influencing whether they self-renew or differentiate into specific cell lineages. The Wnt pathway is particularly important in the context of HFSC development, where Wnt signaling needs to be attenuated to allow HFSC niche establishment. Further, the Wnt pathway is important during homeostatic tissue maintenance in the context of hair follicle cycling, where Wnt signaling is known to play a role in the activation of hair follicle stem cells during the anagen (growth) phase.

It is interesting to note that ceramide availability was shown to regulate Wnt signaling in Drosophila through strong effects on recycling endocytosis of the receptors (Pepperl *et al*, 2013). The Drosophila study implicates Wnt signaling as particularly sensitive to ceramide levels. Given the critical role of Wnt signaling for HFSC differentiation, we postulate that a stem cell-specific ceramide profile is required to accurately fine tune Wnt signaling levels in these cells. While the mechanistic details of how altered ceramide and sphingomyelin profiles lead to misregulated Wnt signaling is not known, it is well known that sphingolipids and ceramides are central to membrane morphology as well as protein sorting, which are critical for Wnt activity (Castanon *et al*, 2020; Pepperl *et al*., 2013; Simons & Ikonen, 1997). High resolution *in vitro* studies of membrane dynamics are now required to understand the role of CerS4 in regulating the dynamics of signaling protein trafficking.

It is interesting to note that ceramide availability was shown to regulate Wnt signaling in Drosophila through strong effects on recycling endocytosis of the receptors (Pepperl *et al*., 2013). The Drosophila study implicates Wnt signaling as particularly sensitive to ceramide levels. Given the critical role of Wnt signaling for HFSC differentiation, we postulate that a stem cell-specific ceramide profile is required to accurately fine tune Wnt signaling levels in these cells. While the mechanistic details of how altered ceramide and sphingomyelin profiles lead to misregulated Wnt signaling is not known, it is well known that sphingolipids and ceramides are central to membrane morphology as well as protein sorting, which are critical for Wnt activity (Harayama & Riezman, 2018; Pinot *et al*, 2014; van Meer *et al*, 2008). High resolution *in vitro* studies of membrane dynamics are now required to understand the role of CerS4 in regulating the dynamics of signaling protein trafficking.

We had previously reported that deletion of CerS4 led to an eczema-like phenotype but the molecular and cellular mechanisms were unclear (Peters *et al*., 2020). We now find that CerS4 expression is restricted to the hair follicle, and the inability of establishment of the HFSC niche is the first phenotype to be detected in the CerS4-deficient mice. Intriguingly, however, the mice gradually develop eczema, implicating the HFSC niche as a key component of the epidermal barrier. Ultrastructural analyses have demonstrated that a large fraction of the skin normal flora resides in hair follicle openings, indicating that hair follicles, in addition to the IFE, are mediators of immune tolerance towards the microbiome (Chen *et al*, 2018). It is known that a specialized microenvironment within hair follicles shields the resident stem cells from immune system surveillance and attack, leading to the HFSC barrier being immune-privileged. The immune privilege is attributed to the presence of immunosuppressive “no danger” signals, local immune-regulatory cells, and the physical seclusion provided by the follicular structure (Agudo *et al*, 2018; Christoph *et al*, 2000; Wang *et al*, 2014). In that respect it is interesting to note that the altered stem cell differentiation in CerS4-deficient HFSCs was associated with reduced expression of immuno-modulatory proteins. Additional mechanistic studies are required to understand the impact of impaired HFSC niche establishment on both the initiation and maintenance of immune tolerance, as this is crucial for unraveling drivers of Th2-driven barrier diseases such as AD, allergic reactions and asthma.

## Materials and Methods

### Mice

C57BL/6N mice with floxed CerS4 alleles were generated in cooperation with Taconic Artemis (Cologne, Germany). CerS4 complete knockout mice were generated as described before (Peters *et al*., 2015). To achieve epidermis-specific deletion, CerS4fl/fl mice were crossed to K14-Cre mice (Hafner *et al*., 2004; Peters *et al*., 2015). To achieve sebaceous gland specific deletion, CerS4^fl/fl^ mice were crossed to SCD3-Cre mice (Dahlhoff *et al*., 2016). Experiments were performed with CerS4^fl/fl^K14Cre^+^ (termed: CerS4^epi-/-^), CerS4^fl/fl^SCD3-Cre^-/+^(termed: CerS4^SCD3-/+^) and CerS4^fl/fl^K14Cre^-^ and CerS4^fl/fl^SCD3-Cre^-/-^ (termed: control) male and female littermates at the indicated postnatal days (P).

### scRNA-sequencing

Back skin of P19 female mice (2 mice / genotype) were dissected and single cell suspensions were generated by digesting cells in a mixture of DNAse I (40 µg/ml), Liberase (0.125mg/ml; Roche Cat.NO. 05401119001 and Papain (30 U/ml Worthington Cat. NO. LK003150) in S- MEM (Gibco) for 60 min at 37 °C. After washing with ice-cold S-MEM, single cells were counted using Luna-II automated cell counter (Logos Biosystems), labeled with Mouse Immune Single-Cell Multiplexing Kit (BD) and loaded on a microwell cartridge of the BD Rhapsody Express system (BD) following the manufacturer’s instructions. Single cell whole transcriptome analysis libraries were prepared according to the manufacturer’s instructions using BD Rhapsody WTA Reagent kit (BD, 633802) and sequenced on the Illumina NextSeq 500 using High Output Kit v2.5 (150 cycles, Illumina) for 2 × 75 bp paired-end reads with 8 bp single index aiming sequencing depth of >20,000 reads per cell for each sample.

Raw FASTQ reads were quality trimmed using fastp (version 0.23.2 length cutoff 20, quality cutoff 15). Adapter trimming was disabled due to the presence of BD Rhapsody multiplexing sequences. The UMI, complex barcode, and sample tags were extracted and demultiplexed using custom scripts.Reads were mapped to the mouse reference genome version GRCm39, with Gencode annotations vM29, filtered for protein-coding genes and lncRNAs, using STAR version 2.7.10a (Dobin *et al*, 2013) (--soloType Droplet –soloCellFilter None –soloFeatures GeneFull_Ex50pAS –soloCBstart 1 –soloCBlen 27 –soloUMIstart 28 –soloUMIlen 8– soloCBwhitelist rhapsody_whitelist.txt –soloMultiMappers EM).

Raw counts were imported as AnnData (Virshup *et al*, 2021) objects. We removed low complexity barcodes with the knee plot method (<1200 counts per cell), and further filtered out cells with a high mitochondrial mRNA percentage (>20%). Doublets were predicted with scrublet (Wolock *et al*, 2019). Finally, each sample’s gene expression matrix was normalized using scran (1.22.1, (Lun *et al*, 2016)) with Leiden clustering (Traag *et al*, 2019) input at resolution 0.5. At this stage samples were merged with scanpy (1.9.1, (Wolf *et al*, 2018)). For 2D embedding, the expression matrix was subset to the 3,000 most highly variable genes (sc.pp.highly_variable_genes, flavor “seurat”). The top 100 principal components (PCs) were calculated, and batch-corrected using Harmony (0.0.5, (Korsunsky *et al*, 2019)). The PCs served as basis for k-nearest neighbor calculation (sc.pp.neighbors, n_neighbors=30), which were used as input for UMAP (McInnes *et al*, 2018) layout (sc.tl.umap, min_dist=0.3). Leiden clustering at resolution 0.8 was used to annotate cell clusters based on previously publsihed marker genes (Joost *et al*., 2016).

#### Pseudobulk DE analysis

To compare gene expression between WT and CerS4^epi-/-^ conditions, we performed “pseudobulk” DE analysis (Squair *et al*, 2021) on the scRNA-seq dataset. First, cells were randomly divided into two pseudoreplicates, and raw expression counts of cells belonging to the same pseudoreplicate were aggregated for each gene. Pseudobulk data was imported into R, and differential expression analysis (DEA) was run using the Wald test for GLM coefficients implemented in the DESeq2 package (1.34.7, (Love *et al*, 2014)). P-values were adjusted for multiple testing using independent hypothesis weighting (IHW, 1.22.0, (Ignatiadis *et al*, 2016)).

#### Trajectory and pseudotime analysis

Lineage trees were calculated independently for WT and CerS4^epi-/-^ using SlingShot (2.2.1, (Street *et al*., 2018)), with the starting cluster chosen as IB. Clusters LH and TC were excluded from the analysis. Diffusion pseudotime (DPT, (Haghverdi *et al*, 2016; Wolf *et al*, 2019) was calculated using the scanpy (1.9.1, (Wolf *et al*., 2018)) functions “sc.tl.diffmap” and “sc.tl.dpt” on an AnnData subset of the clusters: IB, OB1, OB2, uHF1, uHF2, and uHF3. The starting cell was chosen from the IB cluster. Pseudotime expression profiles were generated by binning cells according to (DPT) into 200 bins, and plotting the average normalized expression x alongside a 95% confidence interval (+- 1.96 * sd(x)/sqrt(x)). Cluster labels and colors are assigned based on a “majority vote” of cells in each bin.

#### Ligand-receptor interactions

CellphoneDB (2.1.4, (Efremova *et al*., 2020)) was used to calculate ligand-receptor interactions independently on the WT and CerS4^epi-/-^ conditions (“cellphonedb method statistical_analysis --counts-data hgnc_symbol”). Gene symbols were converted to their human orthologs using Ensembl BioMart (Martin *et al*, 2023).

#### Gene expression comparisons and gene set enrichment

We obtained raw RNA counts from <PMID 25840722> and ran differential expression analysis (DEA) using the Wald test for GLM coefficients implemented in the DESeq2 package (1.34.7, (Love *et al*., 2014)) to compare lesional vs. non-lesional samples. P-values were adjusted for multiple testing using independent hypothesis weighting (IHW, 1.22.0, (Ignatiadis *et al*., 2016)).

For gene set enrichment we ranked differential expression results by −log10(P-value) prefixed with the sign of the log-fold change and used GSEApy (0.10.8, (Fang *et al*, 2023)) “prerank” to calculate the enrichment statistics on various gene sets. Proteomics differential expression results were tested for enrichment against the marker gene sets from Supplementary Table 2 in Joost, Zeisel et al. 2016. To compare atopic dermatitis genes to the gene expression changes in CerS4^epi-/-^ vs WT, we selected DE genes from the lesional vs non-lesional comparison in the atopic dermatitis data (see above) using an adjusted P-value cutoff of 0.0001. Mouse gene symbols were then converted to human gene symbols using Ensembl BioMart (Martin *et al*., 2023) orthologs.

### Hair follicle stem cell organoid culture

Hair follicle stem cell organoids were cultured essentially as described previously (Chacon-Martinez *et al*., 2017). Briefly, epidermal progenitors were isolated from back skin of P19 mice by incubating skin pieces in 0.5% trypsin (Gibco) for 50 min at 37°C. After separating the epidermis from the underlying dermis, cells were passed through 70-μm cell strainer (BD Biosciences) and pelleted at 900 rpm for 3 min. For 3D culture, 8 × 10^4^ cells were suspended in 20 μl ice-cold 1:1 mixture of keratinocyte growth medium (MEM Spinner’s modification (Sigma), 5 µg/mL insulin (Sigma), 10 ng/mL EGF (Sigma), 10 µg/mL transferrin (Sigma), 10 µM phosphoethanolamine (Sigma), 10 µM ethanolamine (Sigma), 0.36 µg/mL hydrocortisone (Calbiochem), 2mM glutamine (Gibco), 100 U/mL penicillin and 100 µg/mL streptomycin (Gibco), and 10% chelated fetal calf serum (Gibco), 5 µM Y27632, 20 ng/mL mouse recombinant VEGF, 20 ng/mL human recombinant FGF-2 (all from Miltenyi Biotec)) and growth factor-reduced Matrigel (Corning) that was dispensed as a droplet in 24-well cell culture dishes. The suspension was allowed to solidify for 30 min after which it was overlaid with 500 μl medium. All cultures were incubated at 37°C, 5% CO2. Medium was exchanged the next day after initial seeding and thereafter every second day. Chir99021 (Sigma # SML1046) was added where indicated.

### Flow cytometry

Single-cell suspensions were prepared from murine back skin as described above or from organoid cultures by mechanical homogenization and incubation in 0.5% Trypsin (Gibco), 0.5 mM EDTA for 10 min at 37°C. Cells were rinsed once with KGM and stained with fluorescently labeled antibodies for 30 min on ice. After two washes with FACS Buffer (2% FCS, 2 mM EDTA, PBS), cells were analyzed in a BD FACSCanto II or sorted in either a BD FACSAria II or a BD FACSAria Fusion. Data were analyzed using FlowJo software version 10.9. Expression of cell surface markers was analyzed on live cells after exclusion of cell doublets and dead cells using 7AAD (eBioscience), 4′, 6-diamidino-2-phenylindole (DAPI, Sigma) or fixable viability dye 405 (eBiosciences). Apoptotic cells were labeled with Alexa Fluor 555-Annexin V (Invitrogen). The following antibodies were used: eFluor660- or FITC- CD34 (clone RAM34, eBioscience), PE-Cy7- or FITC-α6 Integrin (clone GoH3, eBioscience).

Immune cell FACS was performed from ear skin from which cells were isolated using DNAse I 40 µg/ml (Roche, 11284932001) and 20 U/ml Collagenase Type I Worthington (LS0004194) for 90min at 37°C in RPMI1640, followed by gentleMACS tissue dissociation using gentleMACS C tubes and subsequent separation using 70µm cell strainers. After centrifugation 300g, 4°C für 10 min cells were resuspended in 500µl PBS containing 2%FBS.

The following antibodies were used: Fc Block (Invitrogen, 14-0161-86), live/dead eF780 (Invitrogen, 65-0865-14), CD45 FITC (BioLegend, 103108), Gata3 PE (eBioscience, 12- 9966-42), NKp46 eF710 (eBioscience, 46-3351-82), CD4 PE-Cy7 (BioLegend, 100421), Eomes eF660 (eBioscience, 50-4875-82), CD90.2 AF700 (BioLegend 105320), RORgt BV421 (BD Biosciences, 562894), CD25 BV 605 (BD Biosciences, 563061), CD11b super bright 702 (eBioscience 67-0112-82), CD3 Biotin (BioLegend 344820), CD19 Biotin (BioLegend, 115504), TCRb Biotin (BioLegend 109204), TCRgd Biotin (eBioscience, 13- 5711-82), CD5 Biotin (BioLegend (100604), FCeR1 Biotin (BioLegend, 134304), SA V510 anti-Biotin (BioLegend, 405234).

### EdU incorporation

Organoids were grown in the presence or absence of 9.4 μm EdU (Thermo Fischer) for 24 h before analysis. After preparing single-cell suspensions, cells were stained with a fixable viability dye eFluor405 followed by antibody staining before fixation in 2% PFA for 10 min at RT. Cells were subsequently permeabilized in 0.025% Triton X-100, PBS for 10 min, and incubated 30 min in EdU reaction cocktail (100 mM Tris pH 8.5, 1 mM CuSO4, 0.5 μM AlexaFluor 594-Azide (Thermo Fischer), 100 mM ascorbic acid). After two washes with PBS, cells were analyzed by flow cytometry.

### Immunofluorescence and immunohistochemistry

Organoid cultures were rinsed once in PBS, followed by fixation in 2% PFA, PBS for 30 min at RT. Fixed cells were rinsed three times with 100 mM glycine, PBS, then permeabilized and blocked for 2 h at 37°C in 0.3% Triton X-100, 5% BSA, PBS. Cells were stained with primary antibodies in 0.3% Triton X-100, 1% BSA, PBS overnight at RT. Secondary, fluorescent antibodies were used to detect primary antibody binding, and nuclei were visualized with DAPI. Slides were mounted with Elvanol mounting medium. The following antibodies were used: SOX9 (sc-166505, Santa Cruz Biotechnology), Keratin-6 (prb-169P, Convance), CD34 (14-0341, eBioscience). Secondary antibodies anti-mouse AlexaFluor 488 (Invitrogen) and anti-rabbit AlexaFluor 568 (Invitrogen).

Tissue biopsies were fixed (4% PFA), embedded in paraffin and sectioned. Sections were de-paraffinized using a graded alcohol series, blocked in 10% normal goat serum, and incubated with primary antibodies diluted in Dako Antibody Diluent over night at 4°C. Bound primary antibody was detected by incubation with Alexa Fluor 488- or Alexa Fluor 568-conjugated antibodies (Invitrogen). Nuclei were counterstained with 4′, 6-diamidino-2-phenylindole (DAPI, Invitrogen). After washing slides were mounted in Elvanol. The following antibodies were used: Krt14 (Progen #GP-CK14), Krt6 (Covance #prb-169P), Lhx2 (Abcam #ab184337), Lef1 (Cell Signalling #2230S), Nfatc1 (Santa Cruz #sc-7294), and CD45 (eBioscienc2 #12-0451). All fluorescence images were collected by laser scanning confocal microscopy (Zeiss LSM980) controlled by ZEN software (version 3.7), using 40x dry or 63x immersion objective. Images were acquired at room temperature using sequential scanning of frames after which planes were projected as a maximum intensity confocal stack. Images were collected with the same settings for all samples within an experiment. Image processing (linear brightness and contrast enhancement) was performed with Fiji Software version 2.9.0 and Adobe Photoshop CS5.

### RNAScope

Paraffin embedded skin sections were prepared as described above. Sections were labeled using the RNAscope Multiplex Fluorescent Detection Kit v2 (ACDBio, #. 323100). A20zzz probe of murine CerS4 mRNA was designed by the ACD Probe Design Team. The Opal 520 Reagent Pack (Akoya Biosciences, FP1487001KT) was used in a dilution of 1:1000 for the fluorophore step to develop the channel associated with the CerS4 probe. Nuclei were counterstained with 4′, 6-diamidino-2-phenylindole (DAPI, Sigma). Slides were mounted with Elvanol mounting medium.

### qRT-PCR

RNA was isolated using RNeasy Plus Mini Kit (QIAGEN). 500 ng of RNA was subjected to reverse transcription using SuperScript IV VILO Master Mix (Thermo) following the manufacturer’s protocol. PCR was performed with Scientific QuantStudio 5 and 7 Real-Time PCR System (Thermo Fisher) using the DyNAmo Color Flash SYBR Green Mix (Thermo Fisher). Gene expression was quantified using the ΔΔCt method using normalization to GAPDH. Primer sequences were as previously reported (Peters *et al*., 2015).

### Proteomics

Organoid pellets from 4 mice/genotype were lysed and digested using the Preomics iST kit (P.O.00001) according to the manufacturer’s instructions, except that the volume of the LYSE buffer was increased to 70 µl to completely solubilize the organoids. Following peptide elution and lyophilization, peptides were then subjected to TMT labeling in EPPS buffer (20 µl, 200 mM) using TMT labels 127N – 130C (42 µg peptide / replicate and channel; 200 µg labeling reagent). After mixing of the eight channels in an equal ratio, the mixture was desalted on Empore 3M cartridges followed by phosphopeptide enrichment on Fe-NTA spin columns (Thermo Scientific; order # A32992; data not shown). Unbound peptides were lyophilized and offline fractionated by high pH reversed phase HPLC (YMC Triart C18 column, 250 x 4.6 mm ID) into 60 fractions for the subsequent measurements of the total proteomes. All fractions were further desalted on C18 Stage tips prior to mass spectrometry analysis.

Nanoflow LC-MS/MS analysis was carried out as previously published (Bekker-Jensen *et al*, 2017). Briefly, peptide samples were reversed-phase separated on a fused silica capillary column (length 25 cm; ID 75 μm; Dr. Maisch ReproSil-Pur C18-AQ, 1.9 μm) using an Easy nLC 1200 nanoflow system that was online coupled via a Nanospray Flex ion source to an Orbitrap HF mass spectrometer (Thermo Scientific). Bound peptides of each fraction were eluted using short gradients from 7 – 45% B (80% ACN, 0.1% formic acid) in 30 min, followed by a washout at 90% B, before returning again to starting conditions (total runtime 38min). The mass spectrometer was operated in the positive ion mode, switching in a data-dependent fashion between survey scans in the orbitrap (mass range m/z = 350-1400; resolution R=60000; target value = 3E6; max IT 100 ms) and MS/MS acquisition (HCD) of the 20 most intense ion peaks detected (resolution R = 15.000; target value = 1E5; max IT = 15msec; NCE = 27). Dynamic exclusion was enabled and set to 30 s.

Raw MS data were processed using MaxQuant (v. 2.1.4.0) with the built-in Andromeda search engine. Tandem mass spectra were searched against the mouse uniprotKB database (UP000000589_10090.fasta; version from 01/2022) concatenated with reversed sequence versions of all entries and also containing common contaminants. Carbamidomethylation on cysteine residues was set as fixed modification for the search, while oxidation at methionine, acetylation of protein N-termini and deamidation on aspargine and glutamine were set as variable modifications. Trypsin was defined as the digesting enzyme, allowing a maximum of two missed cleavages and requiring a minimum length of 6 amino acids. The maximum allowed mass deviation was 20 ppm for MS and 0.5 Da for MS/MS scans. The match between run function was enabled. Protein groups were regarded as being unequivocally identified with a false discovery rate (FDR) of 1% for both the peptide and protein identifications.

Data transformation and evaluation were performed using Perseus software (version 1.6.15.0). Proteins identified by a single modified site only, common lab contaminants as well as proteins containing reverse sequences derived from the decoy database search were removed from the dataset prior to any further analysis. For quantification, TMT reporter intensity values were first log2 transformed, demanding valid values in all of the replicates in both experimental groups. This step reduced the number of quantifiable protein groups from 6803 to 6609, however, at the same time imputation of missing values was avoided. Reporter intensities were quantile normalized and significant differences between WT controls and KO samples were determined using a Student’s t-test with a permutation-based FDR of 0.05 set as cut-off. Only proteins, which showed an additional > 2-fold difference were considered for further evaluation.

### Lipidomics

Levels of ceramides and sphingomyelins were determined by Liquid Chromatography coupled to Electrospray Ionization Tandem Mass Spectrometry (LC-ESI-MS/MS). Organoids and FACS purified cells were homogenized in Milli-Q water using the Precellys 24 homogenizer (Peqlab) at 6,500 rpm for 30 s. The protein content of the homogenate was routinely determined using bicinchoninic acid. Lipid extraction, alkaline hydrolysis of glycerolipids and LC-ESI-MS/MS analysis of ceramides and sphingomyelins were performed as previously described (Hammerschmidt *et al*, 2023).

### Statistics and reproducibility

Statistical analyses were performed using GraphPad Prism software (GraphPad, version 10). Statistical significance was determined by the specific tests indicated in the corresponding figure legends. Only 2-tailed tests were used. In all cases where a test for normally distributed data was used, normal distribution was confirmed with the Kolmogorov–Smirnov test (α = 0.05). All experiments presented in the manuscript were repeated at least in 3 independent biological replicates.

### Data availability

All data supporting the findings of this study are available from the corresponding author on request. Single cell sequencing data are available at GEO (GSE252821). Proteomics data have been deposited to the ProteomeXchange Consortium through the PRIDE partner repository (Perez-Riverol *et al*, 2022), dataset ID: PXD047573.

## Acknowledgements

We thank Anu Luoto and Sandra Heising for technical assistance, the Max Planck Institute Sequencing Core Facility for support with sequencing, BioOptics and Biomedicum Helsinki Imaging Unit for imaging support and HiLIFE Laboratory Animal Centre Core Facility, University of Helsinki, and Max Planck Institute Animal facility for support with animal experiments. This work was supported by the Max Planck Society, Sigrid Juselius Foundation, Helsinki Institute of Life Science, European Research Council (ERC) under the European Union’s Horizon 2020 research and innovation programme (grant agreement 770877 - STEMpop), and Academy of Finland Center of Excellence BarrierForce (all to SAW). FP is supported by the Walter Benjamin Fellowship from the German Research Foundation (DFG; Project number 455963994).

## Author contribution

FP designed experiments, performed most of the experiments, and analyzed data, SB performed and analyzed the lipidomics experiments, KK analyzed sequencing data, HCAD performed and analyzed proteomics data, JH performed SCD3Cre work, DL and EvSB performed immune cell analyses, MK and CM provided conceptual advice and general support, SAW supervised the study, designed experiments and analyzed data. FP and SAW wrote the manuscript with input from all other authors.

## Supplementary Figures

**Supplementary Figure 1.**
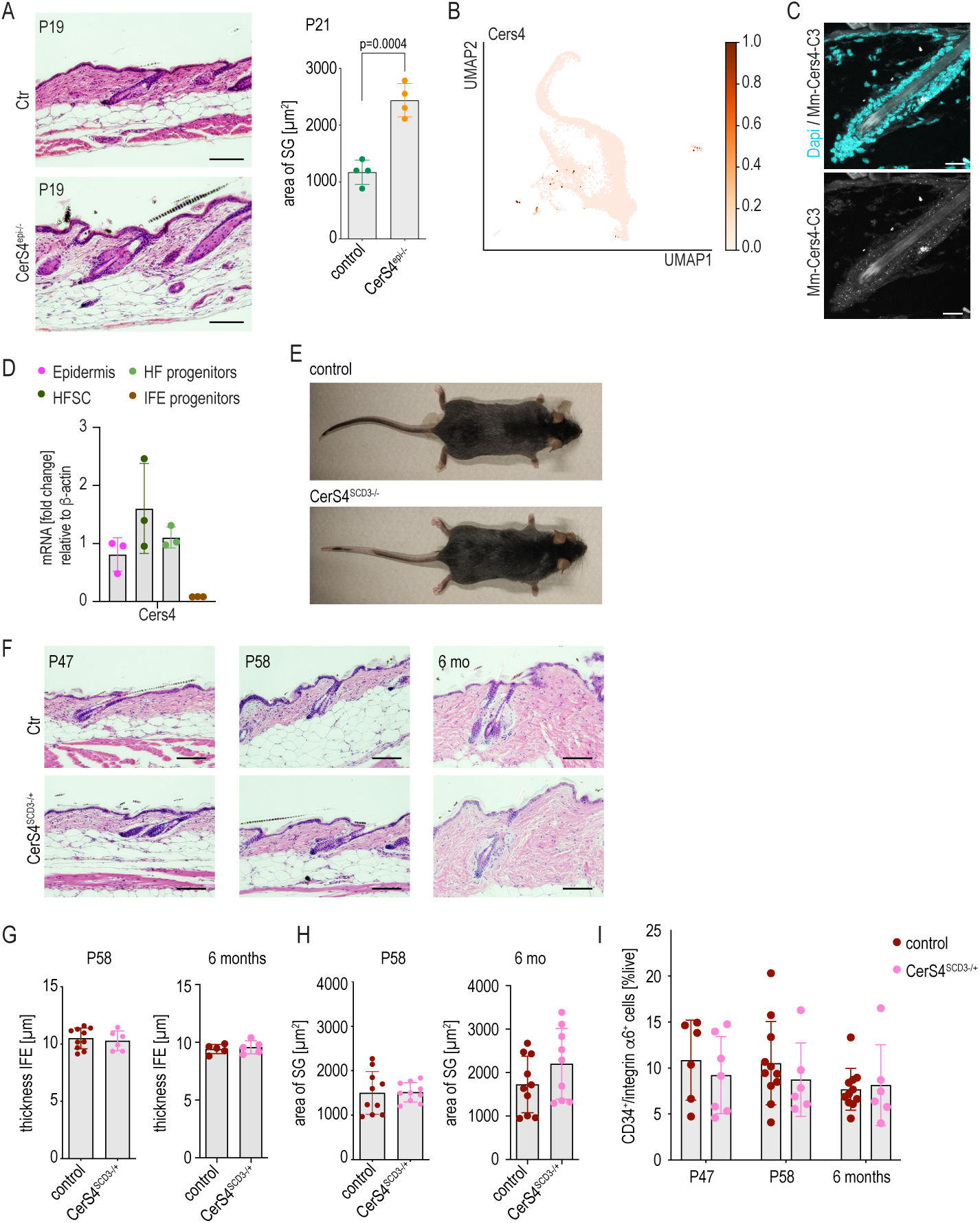
**A.** H/E-stained back skin sections of control and CerS4^epi-/-^ back skin at P19 showing enlarged SGs but otherwise normal HF architecture. Scale bars 100 µm. Quantification of the size of sebaceous glands measured from H/E-stained back skin sections (n=4 mice/genotype; unpaired t-test). **B.** UMAP projection of CerS4 expressing cells in scRNAseq data of control mice. **C.** In situ hybridization of CerS4 mRNA expression in back skin sections of control mice. Scale bars 20µm. **D** QRT-PCR analysis of control FACS sorted HFSCs, HF progenitors and IFE progenitor cells and total epidermis (n=3). **E** Macroscopic images of control and CerS4^SCD3+/-^ mice. **F.** H/E-stained back skin sections of control and CerS4^SCD3+/-^ back skin at P47, P58 and 6 months. Scale bars 100 µm. **G.** Quantification of the thickness of the IFE at P 58 (n=10 control, 6 CerS4^SCD3+/-^ mice) and 6 months (n= 5 mice/genotype) from back skin sections. **H.** Quantification of the size of sebaceous glands (SG) at P 58 (n= 10 mice/genotype) and 6 months (n=10 control, 9 CerS4^SCD3+/-^ mice) from back skin sections. **I.** FACS analysis of CD34^+^ integrin α6^+^ HFSCs from P47 (n=6 control, 7 CerS4^epi-/-^ mice), P58 (n=11 control, 6 CerS4^epi-/-^ mice) and 6 mo old mice (n=11 control, 6 CerS4^epi-/-^ mice).

**Supplementary Figure 2.**
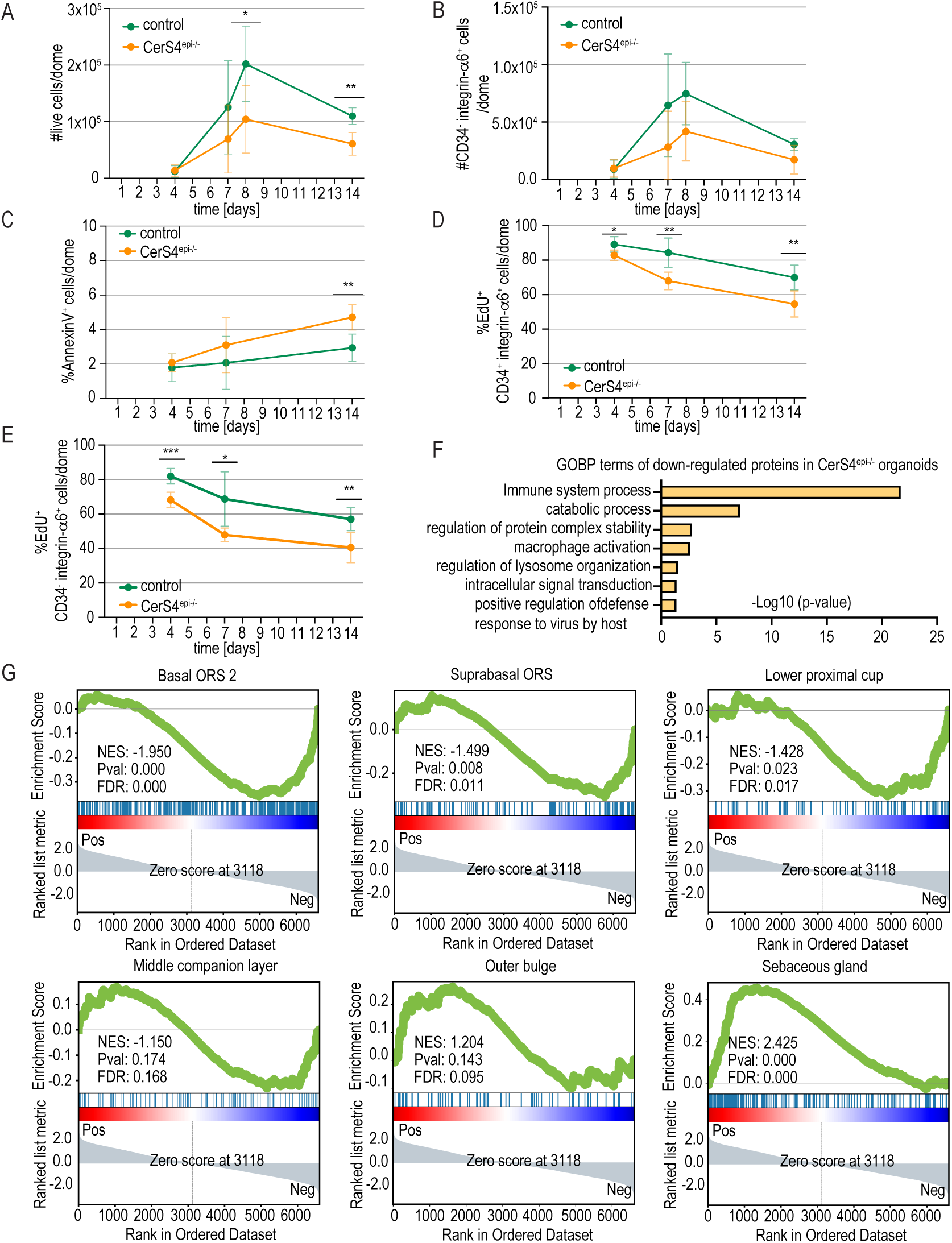
**A, B.** FACS-based quantification of live cells (A) and CD34^-^ integrin α6^+^ non-HFSCs (B) from control and CerS4^epi-/-^ organoids cultured for indicated times (n=4 mice/genotype for 14d, 6 mice/genotype for all other time points; mean ±SD; unpaired t-test). **C.** FACS-based quantification of Annexin-positive cells of control and CerS4^epi-/-^ organoids cultured for indicated time points (n=6 mice/genotype; mean ±SD; unpaired t-test). **D, E.** FACS-based quantification of EdU-positive live cells in CD34^+^ integrin α6^+^ HFSCs (D) and CD34^-^ integrin α6^+^ non-HFSCs (E) after a 24h chase in organoids cultured for indicated time points (n=6 mice/genotype; mean ±SD; unpaired t-test). **F.** GO-term enrichment analysis from proteins downregulated in CerS4^epi-/-^ organoids. **G.** GSEA of known markers of distinct skin hair follicle progenitor cell linages (Joost *et al*., 2016) from differentially expressed proteins in control and CerS4^epi-/-^ organoids.

**Supplementary Figure 3.**
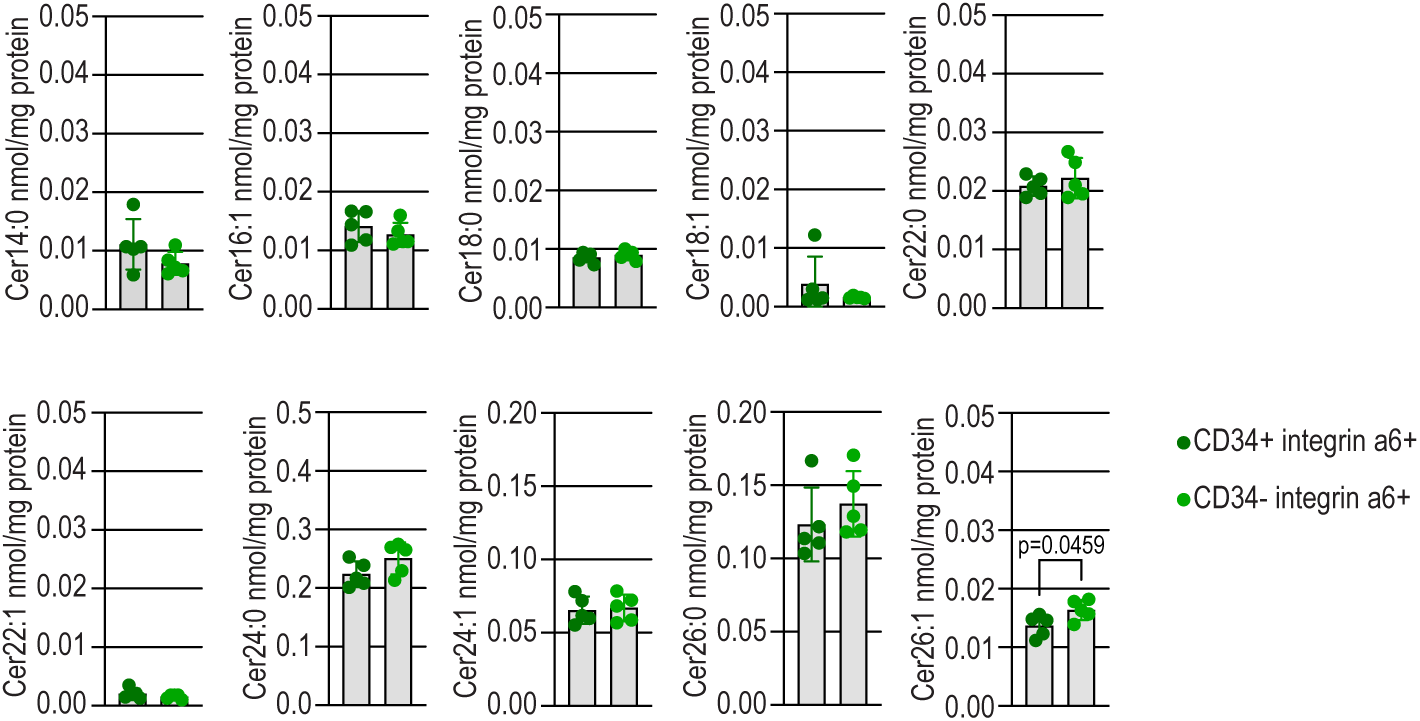
Ceramide levels in FACS purified CD34^+^ integrin α6^+^ HFSCs and CD34^-^ integrin α6^+^ non-HFSCs determined by quantitative LC-ESI-MS/MS analysis (n=5 mice/genotype; ratio paired t-test).

**Supplementary Figure 4.**
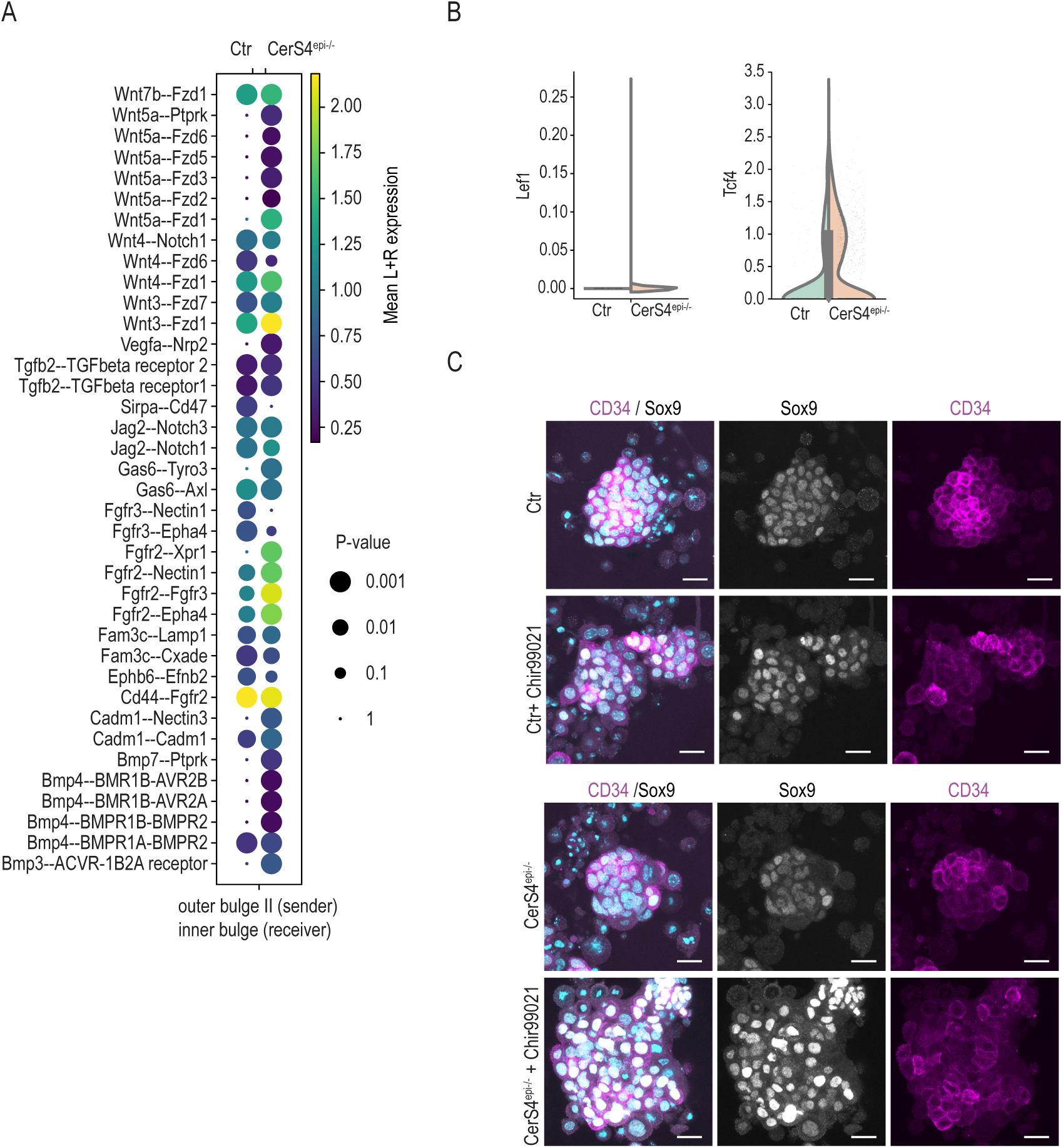
**A.** Receptor-ligand interaction analyses (CellPhoneDB) from scRNAseq data of control and CerS4^epi-/-^ skin. Communication from the outer bulge cells to the inner bulge is illustrated. **B.** Gene expression of Lef1 and Tcf4 in stem and progenitor cell in the outer bulge stem cell compartment from scRNAseq in control and CerS4^epi-/-^ skin. **C.** Representative images of control and CerS4^epi-/-^ organoids treated with Chir99021 and stained for Sox9 (grey) and Keratin 14 (magenta). Scale bars 20µm.

**Supplementary Figure 5.**
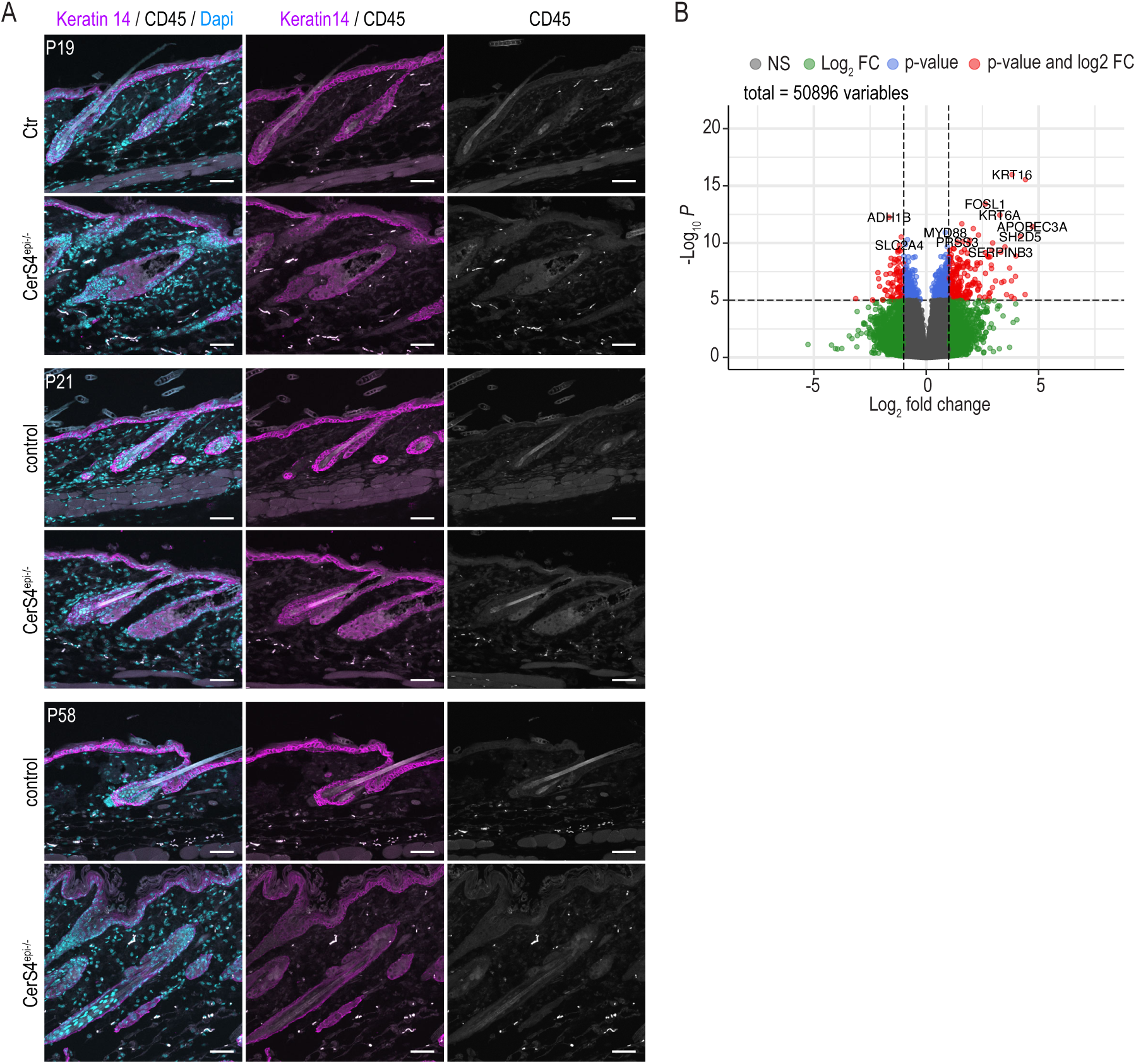
**A.** Representative images of P19, P21 and P58 control and CerS4^epi-/-^ back skin sections stained for CD45 (grey) and Keratin-14 (magenta). Scale bars 50µm. **B** Differential gene expression of 20 patients diagnosed with atopic dermatitis. Positive values indicate genes that are up-regulated in lesional skin and negative values indicate genes that are up-regulated in non-lesional skin from (Federico *et al*., 2020).

## References

1. Agudo J, Park ES, Rose SA, Alibo E, Sweeney R, Dhainaut M, Kobayashi KS, Sachidanandam R, Baccarini A, Merad M et al (2018) Quiescent Tissue Stem Cells Evade Immune Surveillance. Immunity 48: 271–285.e275

2. Bekker-Jensen DB, Kelstrup CD, Batth TS, Larsen SC, Haldrup C, Bramsen JB, Sørensen KD, Høyer S, Ørntoft TF, Andersen CL et al (2017) An Optimized Shotgun Strategy for the Rapid Generation of Comprehensive Human Proteomes. Cell Syst 4: 587–599.e584

3. Bieberich E, MacKinnon S, Silva J, Noggle S, Condie BG (2003) Regulation of cell death in mitotic neural progenitor cells by asymmetric distribution of prostate apoptosis response 4 (PAR-4) and simultaneous elevation of endogenous ceramide. J Cell Biol 162: 469–479

4. Bird CH, Christensen ME, Mangan MS, Prakash MD, Sedelies KA, Smyth MJ, Harper I, Waterhouse NJ, Bird PI (2014) The granzyme B-Serpinb9 axis controls the fate of lymphocytes after lysosomal stress. Cell Death Differ 21: 876–887

5. Blanpain C, Fuchs E (2009) Epidermal homeostasis: a balancing act of stem cells in the skin. Nat Rev Mol Cell Biol 10: 207–217

6. Blanpain C, Fuchs E (2014) Stem cell plasticity. Plasticity of epithelial stem cells in tissue regeneration. Science 344: 1242281

7. Castanon I, Hannich JT, Abrami L, Huber F, Dubois M, Müller M, van der Goot FG, Gonzalez-Gaitan M (2020) Wnt-controlled sphingolipids modulate Anthrax Toxin Receptor palmitoylation to regulate oriented mitosis in zebrafish. Nat Commun 11: 3317

8. Chacon-Martinez CA, Klose M, Niemann C, Glauche I, Wickstrom SA (2017) Hair follicle stem cell cultures reveal self-organizing plasticity of stem cells and their progeny. The EMBO journal 36: 151–164

9. Chacón-Martínez CA, Koester J, Wickström SA (2018) Signaling in the stem cell niche: regulating cell fate, function and plasticity. Development 145

10. Chen YE, Fischbach MA, Belkaid Y (2018) Skin microbiota-host interactions. Nature 553: 427–436

11. Christoph T, Müller-Röver S, Audring H, Tobin DJ, Hermes B, Cotsarelis G, Rückert R, Paus R (2000) The human hair follicle immune system: cellular composition and immune privilege. Br J Dermatol 142: 862–873

12. Dahlhoff M, Camera E, Schafer M, Emrich D, Riethmacher D, Foster A, Paus R, Schneider MR (2016) Sebaceous lipids are essential for water repulsion, protection against UVB-induced apoptosis and ocular integrity in mice. Development 143: 1823–1831

13. Daszczuk P, Mazurek P, Pieczonka TD, Olczak A, Boryń Ł, Kobielak K (2020) An Intrinsic Oscillation of Gene Networks Inside Hair Follicle Stem Cells: An Additional Layer That Can Modulate Hair Stem Cell Activities. Front Cell Dev Biol 8: 595178

14. Dobin A, Davis CA, Schlesinger F, Drenkow J, Zaleski C, Jha S, Batut P, Chaisson M, Gingeras TR (2013) STAR: ultrafast universal RNA-seq aligner. Bioinformatics 29: 15–21

15. Ebel P, Imgrund S, Vom Dorp K, Hofmann K, Maier H, Drake H, Degen J, Dormann P, Eckhardt M, Franz T et al (2014) Ceramide Synthase 4 deficiency in mice causes lipid alterations in sebum and results in alopecia. Biochem J

16. Eckl KM, Tidhar R, Thiele H, Oji V, Hausser I, Brodesser S, Preil ML, Onal-Akan A, Stock F, Muller D et al (2013) Impaired epidermal ceramide synthesis causes autosomal recessive congenital ichthyosis and reveals the importance of ceramide acyl chain length. J Invest Dermatol 133: 2202–2211

17. Efremova M, Vento-Tormo M, Teichmann SA, Vento-Tormo R (2020) CellPhoneDB: inferring cell-cell communication from combined expression of multi-subunit ligand-receptor complexes. Nat Protoc 15: 1484–1506

18. Eyerich K, Eyerich S, Biedermann T (2015) The Multi-Modal Immune Pathogenesis of Atopic Eczema. Trends Immunol 36: 788–801

19. Fang Z, Liu X, Peltz G (2023) GSEApy: a comprehensive package for performing gene set enrichment analysis in Python. Bioinformatics 39

20. Federico A, Hautanen V, Christian N, Kremer A, Serra A, Greco D (2020) Manually curated and harmonised transcriptomics datasets of psoriasis and atopic dermatitis patients. Sci Data 7: 343

21. Folgueras AR, Guo X, Pasolli HA, Stokes N, Polak L, Zheng D, Fuchs E (2013) Architectural niche organization by LHX2 is linked to hair follicle stem cell function. Cell Stem Cell 13: 314–327

22. Furio L, Pampalakis G, Michael IP, Nagy A, Sotiropoulou G, Hovnanian A (2015) KLK5 Inactivation Reverses Cutaneous Hallmarks of Netherton Syndrome. PLoS Genet 11: e1005389

23. Gonzales KAU, Fuchs E (2017) Skin and Its Regenerative Powers: An Alliance between Stem Cells and Their Niche. Dev Cell 43: 387–401

24. Hafner M, Wenk J, Nenci A, Pasparakis M, Scharffetter-Kochanek K, Smyth N, Peters T, Kess D, Holtkotter O, Shephard P et al (2004) Keratin 14 Cre transgenic mice authenticate keratin 14 as an oocyte-expressed protein. Genesis 38: 176–181

25. Haghverdi L, Buttner M, Wolf FA, Buettner F, Theis FJ (2016) Diffusion pseudotime robustly reconstructs lineage branching. Nature methods 13: 845–848

26. Hammerschmidt P, Steculorum SM, Bandet CL, Del Río-Martín A, Steuernagel L, Kohlhaas V, Feldmann M, Varela L, Majcher A, Quatorze Correia M et al (2023) CerS6-dependent ceramide synthesis in hypothalamic neurons promotes ER/mitochondrial stress and impairs glucose homeostasis in obese mice. Nat Commun 14: 7824

27. Harayama T, Riezman H (2018) Understanding the diversity of membrane lipid composition. Nature reviews Molecular cell biology 19: 281–296

28. He Q, Wang G, Wakade S, Dasgupta S, Dinkins M, Kong JN, Spassieva SD, Bieberich E (2014) Primary cilia in stem cells and neural progenitors are regulated by neutral sphingomyelinase 2 and ceramide. Mol Biol Cell 25: 1715–1729

29. Horsley V, Aliprantis AO, Polak L, Glimcher LH, Fuchs E (2008) NFATc1 balances quiescence and proliferation of skin stem cells. Cell 132: 299–310

30. Hsu YC, Li L, Fuchs E (2014) Emerging interactions between skin stem cells and their niches. Nat Med 20: 847–856

31. Hsu YC, Pasolli HA, Fuchs E (2011) Dynamics between stem cells, niche, and progeny in the hair follicle. Cell 144: 92–105

32. Ignatiadis N, Klaus B, Zaugg JB, Huber W (2016) Data-driven hypothesis weighting increases detection power in genome-scale multiple testing. Nature methods 13: 577–580

33. Jennemann R, Rabionet M, Gorgas K, Epstein S, Dalpke A, Rothermel U, Bayerle A, van der Hoeven F, Imgrund S, Kirsch J et al (2012) Loss of ceramide synthase 3 causes lethal skin barrier disruption. Hum Mol Genet 21: 586–608

34. Joost S, Zeisel A, Jacob T, Sun X, La Manno G, Lonnerberg P, Linnarsson S, Kasper M (2016) Single-Cell Transcriptomics Reveals that Differentiation and Spatial Signatures Shape Epidermal and Hair Follicle Heterogeneity. Cell Syst 3: 221–237 e229

35. Kato T, Liu N, Morinaga H, Asakawa K, Muraguchi T, Muroyama Y, Shimokawa M, Matsumura H, Nishimori Y, Tan LJ et al (2021) Dynamic stem cell selection safeguards the genomic integrity of the epidermis. Dev Cell 56: 3309–3320.e3305

36. Kim CS, Ding X, Allmeroth K, Biggs LC, Kolenc OI, L’Hoest N, Chacón-Martínez CA, Edlich-Muth C, Giavalisco P, Quinn KP et al (2020) Glutamine Metabolism Controls Stem Cell Fate Reversibility and Long-Term Maintenance in the Hair Follicle. Cell Metab 32: 629–642.e628

37. Korsunsky I, Millard N, Fan J, Slowikowski K, Zhang F, Wei K, Baglaenko Y, Brenner M, Loh PR, Raychaudhuri S (2019) Fast, sensitive and accurate integration of single-cell data with Harmony. Nature methods 16: 1289–1296

38. Levy M, Futerman AH (2010) Mammalian ceramide synthases. IUBMB Life 62: 347–356

39. Love MI, Huber W, Anders S (2014) Moderated estimation of fold change and dispersion for RNA-seq data with DESeq2. Genome Biol 15: 550

40. Lun AT, Bach K, Marioni JC (2016) Pooling across cells to normalize single-cell RNA sequencing data with many zero counts. Genome Biol 17: 75

41. Martin FJ, Amode MR, Aneja A, Austine-Orimoloye O, Azov AG, Barnes I, Becker A, Bennett R, Berry A, Bhai J et al (2023) Ensembl 2023. Nucleic acids research 51: D933–D941

42. McInnes L, Healy J, Melville J (2018) UMAP: Uniform Manifold Approximation and Projection for Dimension Reduction. arXiv 1802: 03426

43. Mullen TD, Spassieva S, Jenkins RW, Kitatani K, Bielawski J, Hannun YA, Obeid LM (2011) Selective knockdown of ceramide synthases reveals complex interregulation of sphingolipid metabolism. J Lipid Res 52: 68–77

44. Neill DR, Wong SH, Bellosi A, Flynn RJ, Daly M, Langford TK, Bucks C, Kane CM, Fallon PG, Pannell R et al (2010) Nuocytes represent a new innate effector leukocyte that mediates type-2 immunity. Nature 464: 1367–1370

45. Paternoster L, Standl M, Waage J, Baurecht H, Hotze M, Strachan DP, Curtin JA, Bønnelykke K, Tian C, Takahashi A et al (2015) Multi-ancestry genome-wide association study of 21,000 cases and 95,000 controls identifies new risk loci for atopic dermatitis. Nat Genet 47: 1449–1456

46. Pepperl J, Reim G, Luthi U, Kaech A, Hausmann G, Basler K (2013) Sphingolipid depletion impairs endocytic traffic and inhibits Wingless signaling. Mech Dev 130: 493–505

47. Perez-Riverol Y, Bai J, Bandla C, García-Seisdedos D, Hewapathirana S, Kamatchinathan S, Kundu DJ, Prakash A, Frericks-Zipper A, Eisenacher M et al (2022) The PRIDE database resources in 2022: a hub for mass spectrometry-based proteomics evidences. Nucleic Acids Res 50: D543–D552

48. Perng YC, Lenschow DJ (2018) ISG15 in antiviral immunity and beyond. Nat Rev Microbiol 16: 423–439

49. Peters F, Tellkamp F, Brodesser S, Wachsmuth E, Tosetti B, Karow U, Bloch W, Utermohlen O, Kronke M, Niessen CM (2020) Murine Epidermal Ceramide Synthase 4 Is a Key Regulator of Skin Barrier Homeostasis. J Invest Dermatol

50. Peters F, Vorhagen S, Brodesser S, Jakobshagen K, Brüning JC, Niessen CM, Krönke M (2015) Ceramide synthase 4 regulates stem cell homeostasis and hair follicle cycling. J Invest Dermatol 135: 1501–1509

51. Pinot M, Vanni S, Pagnotta S, Lacas-Gervais S, Payet LA, Ferreira T, Gautier R, Goud B, Antonny B, Barelli H (2014) Lipid cell biology. Polyunsaturated phospholipids facilitate membrane deformation and fission by endocytic proteins. Science 345: 693–697

52. Radner FP, Marrakchi S, Kirchmeier P, Kim GJ, Ribierre F, Kamoun B, Abid L, Leipoldt M, Turki H, Schempp W et al (2013) Mutations in CERS3 cause autosomal recessive congenital ichthyosis in humans. PLoS Genet 9: e1003536

53. Salimi M, Barlow JL, Saunders SP, Xue L, Gutowska-Owsiak D, Wang X, Huang LC, Johnson D, Scanlon ST, McKenzie AN et al (2013) A role for IL-25 and IL-33-driven type-2 innate lymphoid cells in atopic dermatitis. J Exp Med 210: 2939–2950

54. Schneider MR, Schmidt-Ullrich R, Paus R (2009) The hair follicle as a dynamic miniorgan. Curr Biol 19: R132–142

55. Simons K, Ikonen E (1997) Functional rafts in cell membranes. Nature 387: 569–572

56. Squair JW, Gautier M, Kathe C, Anderson MA, James ND, Hutson TH, Hudelle R, Qaiser T, Matson KJE, Barraud Q et al (2021) Confronting false discoveries in single-cell differential expression. Nature communications 12: 5692

57. Street K, Risso D, Fletcher RB, Das D, Ngai J, Yosef N, Purdom E, Dudoit S (2018) Slingshot: cell lineage and pseudotime inference for single-cell transcriptomics. BMC Genomics 19: 477

58. Subramanian A, Tamayo P, Mootha VK, Mukherjee S, Ebert BL, Gillette MA, Paulovich A, Pomeroy SL, Golub TR, Lander ES et al (2005) Gene set enrichment analysis: a knowledge-based approach for interpreting genome-wide expression profiles. Proc Natl Acad Sci U S A 102: 15545–15550

59. Tidhar R, Zelnik ID, Volpert G, Ben-Dor S, Kelly S, Merrill AH, Futerman AH (2018) Eleven residues determine the acyl chain specificity of ceramide synthases. J Biol Chem 293: 9912–9921

60. Traag VA, Waltman L, van Eck NJ (2019) From Louvain to Leiden: guaranteeing well-connected communities. Scientific reports 9: 5233

61. Tretina K, Park ES, Maminska A, MacMicking JD (2019) Interferon-induced guanylate-binding proteins: Guardians of host defense in health and disease. J Exp Med 216: 482–500

62. Tumbar T, Guasch G, Greco V, Blanpain C, Lowry WE, Rendl M, Fuchs E (2004) Defining the epithelial stem cell niche in skin. Science 303: 359–363

63. van Gastel N, Stegen S, Eelen G, Schoors S, Carlier A, Daniels VW, Baryawno N, Przybylski D, Depypere M, Stiers PJ et al (2020) Lipid availability determines fate of skeletal progenitor cells via SOX9. Nature 579: 111–117

64. van Meer G, Voelker DR, Feigenson GW (2008) Membrane lipids: where they are and how they behave. Nature reviews Molecular cell biology 9: 112–124

65. Virshup I, Rybakov S, Theis FJ, Angerer P, Wolf FA (2021) anndata: Annotated data. bioRxiv: 2021.2012.2016.473007

66. Vivier E, Artis D, Colonna M, Diefenbach A, Di Santo JP, Eberl G, Koyasu S, Locksley RM, McKenzie ANJ, Mebius RE et al (2018) Innate Lymphoid Cells: 10 Years On. Cell 174: 1054–1066

67. Wang G, Krishnamurthy K, Chiang YW, Dasgupta S, Bieberich E (2008) Regulation of neural progenitor cell motility by ceramide and potential implications for mouse brain development. J Neurochem 106: 718–733

68. Wang X, Marr AK, Breitkopf T, Leung G, Hao J, Wang E, Kwong N, Akhoundsadegh N, Chen L, Mui A et al (2014) Hair follicle mesenchyme-associated PD-L1 regulates T-cell activation induced apoptosis: a potential mechanism of immune privilege. J Invest Dermatol 134: 736–745

69. Weidinger S, Beck LA, Bieber T, Kabashima K, Irvine AD (2018) Atopic dermatitis. Nat Rev Dis Primers 4: 1

70. Weidinger S, Novak N (2016) Atopic dermatitis. Lancet 387: 1109–1122

71. Wolf FA, Angerer P, Theis FJ (2018) SCANPY: large-scale single-cell gene expression data analysis. Genome Biol 19: 15

72. Wolf FA, Hamey FK, Plass M, Solana J, Dahlin JS, Gottgens B, Rajewsky N, Simon L, Theis FJ (2019) PAGA: graph abstraction reconciles clustering with trajectory inference through a topology preserving map of single cells. Genome Biol 20: 59

73. Wolock SL, Lopez R, Klein AM (2019) Scrublet: Computational Identification of Cell Doublets in Single-Cell Transcriptomic Data. Cell systems 8: 281–291 e289

74. Xu Z, Wang W, Jiang K, Yu Z, Huang H, Wang F, Zhou B, Chen T (2015) Embryonic attenuated Wnt/β-catenin signaling defines niche location and long-term stem cell fate in hair follicle. Elife 4: e10567

75. Yang T, Liang D, Koch PJ, Hohl D, Kheradmand F, Overbeek PA (2004) Epidermal detachment, desmosomal dissociation, and destabilization of corneodesmosin in Spink5-/- mice. Genes Dev 18: 2354–2358

76. Ziegler C, Graf J, Faderl S, Schedlbauer J, Strieder N, Förstl B, Spang R, Bruckmann A, Merkl R, Hombach S et al (2019) The long non-coding RNA LINC00941 and SPRR5 are novel regulators of human epidermal homeostasis. EMBO Rep 20

